# Dinucleotides as simple models of the base stacking-unstacking component of DNA ‘breathing’ mechanisms

**DOI:** 10.1101/2020.10.26.355974

**Authors:** Eric R. Beyerle, Mohammadhasan Dinpajooh, Huiying Ji, Peter H. von Hippel, Andrew H. Marcus, Marina G. Guenza

## Abstract

Regulatory protein access to the DNA duplex ‘interior’ depends on local DNA ‘breathing’ fluctuations, and the most fundamental of these are thermally-driven base stacking-unstacking interactions. The smallest DNA unit that can undergo such transitions is the dinucleotide, whose structural and dynamic properties are dominated by stacking, while the ion condensation, cooperative stacking and inter-base hydrogen-bonding, present in duplex DNA are not involved. We use dApdA to study stacking-unstacking at the dinucleotide level because the fluctuations observed are likely to resemble those of larger DNA molecules, but in the absence of constraints introduced by cooperativity are likely to be more pronounced, and thus more accessible to measurement. We study these fluctuations with a combination of Molecular Dynamics simulations on the microsecond timescale and Markov State Model analyses, and validate our results by calculations of circular dichroism (CD) spectra, with results that agree well with experiments. Our analyses show that the CD spectrum of dApdA is defined by two distinct chiral conformations that correspond, respectively, to a Watson-Crick form and a hybrid form with one base in a Hoogsteen configuration. We find also that ionic structure and water orientation around dApdA play important roles in controlling its breathing fluctuations.

## Introduction

Nucleic acids undergo a variety of local structural fluctuations in discharging their biological functions. These fluctuations (collectively called ‘breathing’) include inter-strand base-pair opening and closing, intra-strand base stacking and unstacking and conformational rearrangements of the sugar-phosphate backbone. ^1,2,3,4,5,6^ Such thermally activated DNA ‘breathing’ fluctuations are thought to represent primary steps in the process by which genome regulatory proteins gain access to the double-stranded (ds) DNA interior.

Understanding thermally driven DNA fluctuations may provide a central key to structural and dynamic interpretation of the interactions between functional and regulatory proteins and their ss- and dsDNA targets during gene expression. However, many of these ‘breathing’ processes, if considered only in duplex DNA, are likely to represent a small fraction of the population of conformations present in duplex DNA at physiological temperatures because of ‘structural cooperativity’ and may thus be hard to resolve even by sensitive spectroscopic and computational techniques. One way of reducing this problem is to focus on elementary systems, such as dinucleotides. These can be considered to represent the ‘fundamental fragments’ of duplex DNA, but also provide a milieu in which the only relevant breathing process is likely to be base stacking and unstacking. As a consequence, these processes can be studied in isolation in these small model systems. In addition, because of the absence of constraints imposed by neighboring and base-paired nucleotides, these stacking-unstacking fluctuations are likely to be present at higher concentrations than in larger duplex DNA molecules and thus also more amenable to study. These considerations have motivated us to reinvestigate the structure and dynamics of dApdA as a model dinucleotide fragment of duplex DNA using modern computational and molecular modeling techniques.

The relative populations of stacked and unstacked bases present in DNA molecules in solution under a variety of environmental conditions have traditionally been studied by absorbance and circular dichroism (CD) experiments. ^7,8^ Initial studies of DNA stacking-unstacking fluctuations focused on dinucleotides in solution.^9,10,11,12,13,14,15^ Dinucleotides, such as dApdA, have been considered to be useful models for some of the basic interactions that control and stabilize local base conformations of dsDNA because – as indicated above – stacking interactions can be examined in these systems while avoiding the complicating features of ion condensation, cooperative stacking and inter-base hydrogen-bonding that are also present and involved in controlling the conformational behavior of long duplex DNA. In addition, homo-dinucleotides, such as dApdA, are more useful than hetero-dinucleotides as model systems for probing conformational rearrangements in these structures because the CD signals from homo-dinucleotides are strengthened by the presence of degenerate exciton coupling effects. Furthermore, dinucleotides may also serve as partial models for deciphering the structure and energetics of some of the more complex elements of biologically important DNA structure, such as the single-stranded (ss) DNA—dsDNA forks and junctions that are essential intermediates in the pathways by which proteins that control genome expression find and interact with their target sites, but in which cooperative interactions and hydrogen bonds between strands are not significantly present.

Base and base-pair interaction free energies have typically been estimated from thermal denaturation studies of DNA oligonucleotides,^16,17^ which showed that among the contributions to the overall interaction free energies of these systems, the free energy of hydrogen bonding between complementary bases and the energetics of configurational and solvent entropy provide only small contributions to the stability of the base paired structures.^14^ Furthermore, base-base stacking, which is the main (enthalpic) contributor to the stability of dinucleotide conformations, appears also to be the dominant component of the overall stability of more complex DNA structures.^10,18,19^ Early studies of dApdA by Schellman, Tinoco and others ^8,10,12,13,14,18^ suggested that the CD spectrum of this dinucleotide in aqueous salt solutions could be represented as the weighted sum of two conformations, one ‘stacked’ and the other ‘unstacked’, with the stacked form likely resembling (in terms of base-base overlap and helical pitch) the Watson-Crick B-form characteristic of duplex DNA. Furthermore, these workers showed that the changes induced in the CD spectrum of this dinucleotide by increasing concentrations of monovalent salt (NaCl) could be attributed to shifts in the relative populations of these same two conformations.

However, these interpretations clearly represented over-simplifications of the actual situation, since we now know that the CD spectrum of a given molecule of this sort must comprise a sum over myriad microstate configurations that simultaneously exist in solution at equilibrium. As a consequence of this complex situation, CD spectra cannot be ‘inverted’ to determine the conformations that contribute uniquely to the overall spectrum. We here address this problem by means of extensive Molecular Dynamics (MD) simulations and a Markov State Model (MSM) analysis,^20,21,22^ thus providing information on the major conformations that participate in the stacking-unstacking equilibria of dApdA, and whose excitonic transitions contribute to the overall CD spectrum.

To this end we performed a set of 2 μs MD simulations of the dApdA dinucleotide in aqueous solvent at increasing monovalent salt concentrations, using the same conditions employed for the initial spectroscopic measurements on dApdA dinucleotide.^15^ From our MD trajectories, each consisting of ~10^7^ microstate configurations, we calculated the CD spectrum by averaging together the contributions from each MD-generated conformation using the standard method^11,12,23,24^ together with an extended dipole model (EDM).^24^ The initial predictions generated by this method showed excellent agreement with experiment.

We next carried out an MSM analysis of our MD trajectories and identified five kinetically stable regions in the free energy landscape, which we refer to as ‘macrostates.’ Each macrostate contains a ‘family’ of conformationally-related microstates, which rapidly interconvert. Transitions between macrostates are kinetically uncoupled, since they cross high energy barriers and thus follow Markovian statistics.^20^ The ensemble of macrostates establishes a structural basis for the interpretation of spectroscopic measurements. By combining MSM analyses with transition path sampling^25,26,27,28,29^ we investigated the kinetic pathways for base unstacking, thus revealing the roles that base flipping appears to play in breathing fluctuations at the dinucleotide level. In addition, we were able to identify one average configuration for each macrostate that served, with sufficient accuracy, to represent the averaged properties of the macrostate. This simplified, five-configuration model retains the important features of the CD spectrum calculated using the full MD statistics and provides a useful minimalistic ensemble for the calculation of CD and potentially other optical spectra using more sophisticated experimental techniques.

Of the five macrostates, three are statistically the most populated, with the CD spectrum being largely determined as the sum of contributions from only two configurational states, consistent with early experimental observations.^10^ While the original studies interpreted those spectra in terms of a single stacked and a single unstacked configuration of dApdA, our analysis shows that, of the two conformational states that contribute significantly to the features of the CD spectra, the most populated corresponds to an ensemble of hybrid dinucleotide conformations that include one base that has flipped into a *syn* conformation, which in dsDNA results in Hoogsteen base-pairing, ^30,31,32,33,34,35,36,37^ and the relatively less populated state corresponds to an ensemble in which both bases of the dinucleotide remain in the canonical *anti* conformation, compatible with right-handed B form (Watson-Crick) base-pairing in dsDNA. The third highly populated dApdA conformation, which is partially unstacked, and contains one *syn* base, does not contribute significantly to the CD signal. However, these results do indicate that conformations compatible with the Hoogsten structure could well play an important role in some types of breathing fluctuations—at least at the dinucleotide level -- thus confirming its possible relevance to biologically important breathing fluctuations in larger DNA molecules as well.^30–37^

Our studies of the orientations and distributions of counterions in aqueous solutions of dApdA have revealed an abrupt structural transition in the positioning and distribution of these ions around the dinucleotide at a NaCl concentration slightly above 1 M, indicating that counterion concentrations are also involved in controlling breathing fluctuations at the dinucleotide level,^19,38^ and likely in larger DNA molecules as well. We show that the above abrupt salt-concentration-dependent transition is correlated with a shift in the equilibria between the three most populated macrostates of the dApdA dinucleotide, and is consistent with early thermal studies of DNA stability at increasing monovalent salt concentration.^39,40,41^ We have shown that this transition is not seen in MD simulations of the isolated phosphate anion in ionic solutions, suggesting that this salt-dependent transition depends also on other (uncharged) elements of the dinucleotide structure.

## Material and Methods

Molecular dynamics (MD) simulations of the dApdA dinucleotide monophosphate in the NPT ensemble were performed in aqueous solution at increasing salt concentrations and analyzed using the Markov state model (MSM) summarized in the appended Supplementary Information (SI).

The theoretical modeling of circular dichroism (CD) Spectra of the dApdA dinucleotide monophosphate was performed using the extended-dipole model (EDM) following the standard CD calculations summarized in the appended Supplementary Information (SI).

## Results

### Structural parameters of dinucleotides

As pointed out above, it has long been known that under physiological salt conditions the adjacent bases of each strand of duplex DNA in aqueous solution adopt helical conformations close to the Watson-Crick B-form, with average inter-base separation *R* ~ 3.5 Å and relative twist angle *φ* ~ 36° (see Fig. 1 for parameter definitions). Spectroscopic studies of small oligonucleotides in solution have examined the various contributions to base stacking stability in duplex and ssDNA – i.e., the effects of hydrophobic bonding, backbone interactions, inter-base hydrogen bonding and cooperativity.^8,18^

**Figure 1.**
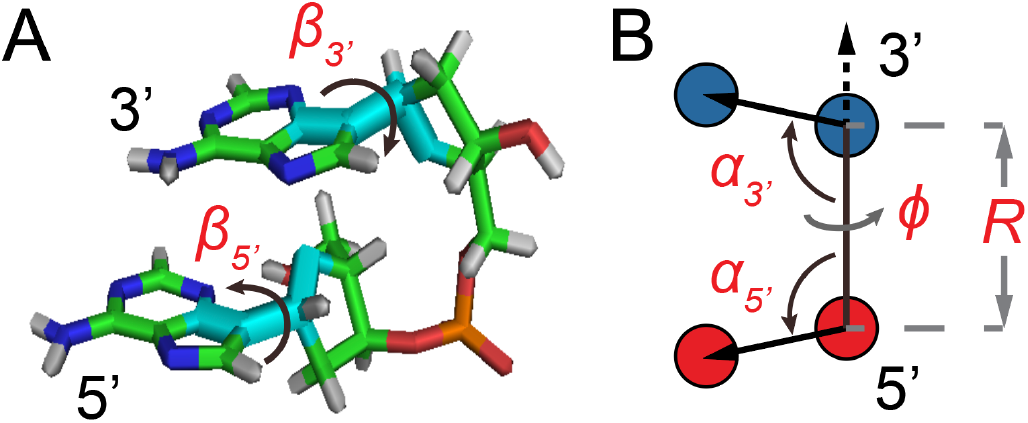
Structural coordinates for the dApdA dinucleotide monophosphate used in this work. (***A***) An atomic scale structure is shown with inter-base roll angles *β*_3’_ and *β*_5’_. (***B***) Virtual atoms are shown with blue and red spheres positioned within the planes of the 5’ and 3’ bases, respectively, with inter-base separation *R*, tilt angles *α*_*S’*_ and *α*_*5’*_, and dihedral twist *φ* (see SI for further details).

### Free energy landscapes as a function of structural parameters and varying salt concentration

Prior studies of the dApdA dinucleotide monophosphate used CD spectroscopy to investigate changes in base conformation as a function of salt concentration, in order to elucidate the roles of the solvent ions in controlling dinucleotide conformation.^8,18^ These studies concluded that the predominant conformation for these truncated ssDNA molecules at physiological salt conditions is a stacked form close to the right-handed Watson-Crick B-form conformation, and that increasing the salt concentration appeared to destabilize this B-form conformation. As we discuss further below, the results of our analyses suggest that the dApdA system is, in fact, more accurately described as an equilibrium distribution of primarily three distinct stacked conformations.

We performed MD simulations of dApdA in aqueous solution at increasing salt concentrations with [NaCl] = 0.1, 0.5, 1.0, 1.05, 1.2 and 1.5 M, as described in the Methods and the SI sections (see Fig. S1 for a snapshot of a sample configuration). For each 2 μs simulation run, we constructed a histogram representing the probability *P*(*R*, *φ*) of finding a configuration at a given value of the structural parameters *R* and *φ* and the related free energy values, *G*(*R*, *φ*) = −*k*_*B*_*T* ln *P*(*R*, *φ*) (see also Fig. S5, which reports the parameters adopted to calculate the FES and perform the MSM analysis, and related discussion in the SI). An example of such a free energy contour diagram plot is shown in Fig. 2*A*. To ensure that the FES represents the system at equilibrium, we show, in Fig. S9 of the SI, the time autocorrelation function of the fluctuations in inter-base separation. Because the function is found to decay in ~10 ns, which compares well with the 2 *μ*s of simulation time, the system can be considered to be in equilibrium.

**Figure 2.**
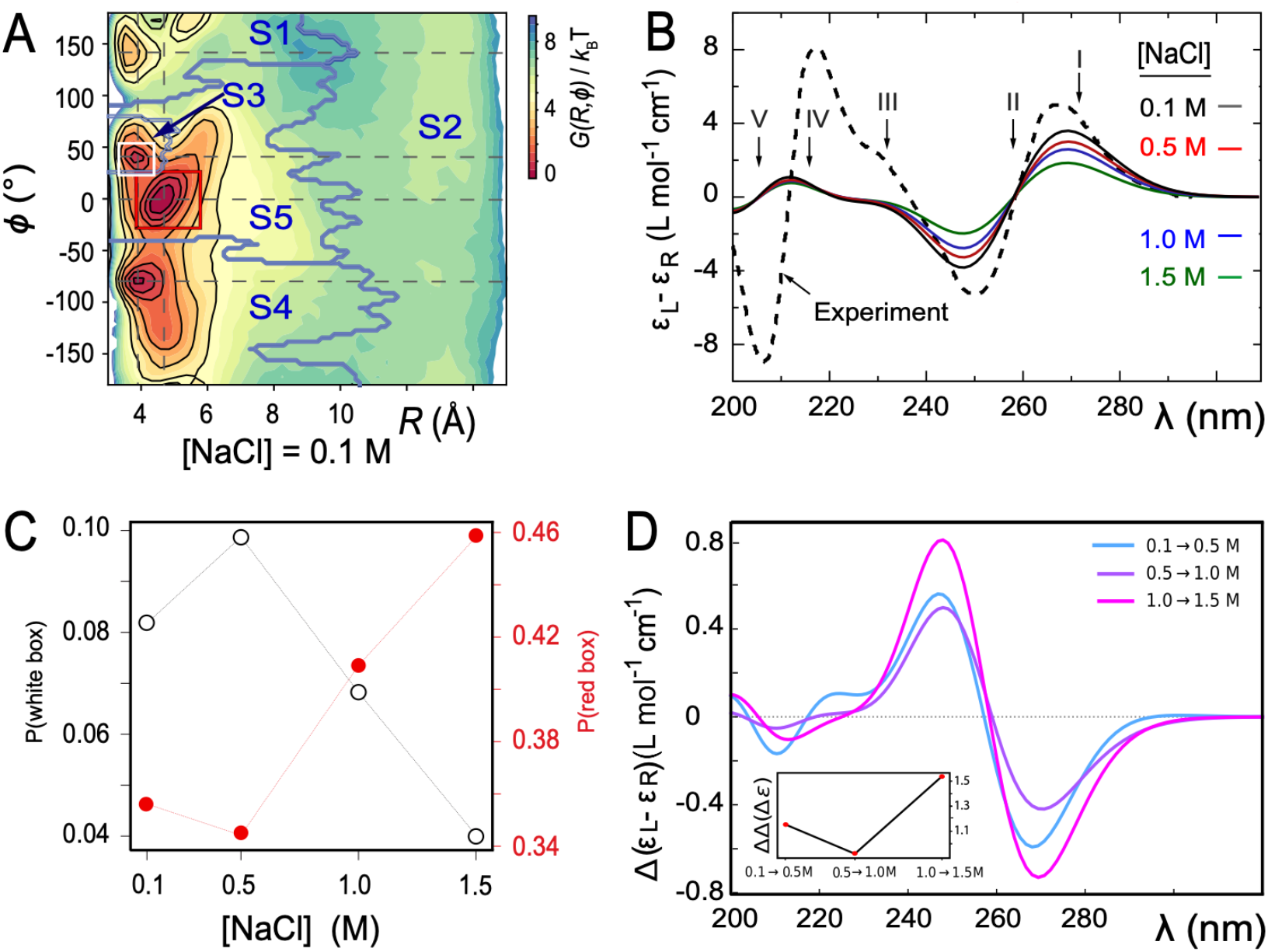
(***A***) Free energy landscape *G*(*R*, *φ*) as a function of the inter-base separation *R* and the dihedral twist angle *φ*, as obtained from 2 μs MD simulations of the dApdA dinucleotide with [NaCl] = 0.1 M. The coordinates corresponding to the canonical (average) B-form conformation (*R* = 3.6 Å and *φ* = 36°) are included in the white square, while the unstacked ‘achiral’ conformations are included in the red square. The five macrostate regions, labeled S1–S5, were identified through the Markov State Modeling procedure (see related section). (***B***) CD spectra of dApdA were determined from 2 μs MD simulations at salt concentrations [NaCl] = 0.1 (black), 0.5 (red), 1.0 (blue) and 1.5 M (green). Differences between the calculated spectra are greater than the error bars (shown as the width of the colored lines), which were determined from the standard error of the mean from five block averages. The experimental CD spectrum (dashed black curve) was taken from Ref. 15. Roman numerals indicate the wavelengths of the electronic transitions of the uncoupled adenine monomers (see Table S1 in the SI), which are used as input for our calculations. (***C***) The local probabilities of the B-like stacked conformation and the unstacked ‘achiral’ conformation was calculated as the sum of states contained within the boundaries defined by the white and red squares, respectively, shown in panel (*A*). These probabilities are shown as a function of salt concentration [NaCl] = 0.1, 0.5, 1.0 and 1.5 M. The relative population of stacked and unstacked conformations changes abruptly around 1 M concentration. (***D***) Differences between the CD spectra at increasing salt concentration: the difference between calculated CD spectra are shown for incremental increases of the salt concentration. The peak-to-peak amplitude of the difference CD spectra decreases dramatically when the concentration is raised above [NaCl] = 1 M. This is reflected by the abrupt change in the peak-to-peak amplitude of the difference CD spectra shown in the inset.

Two additional sets of orientational coordinates per base – the base tilt angles, *α*_*S’*_ and *α*_*S’*_, and the roll angles, *β*_3’_ and *β*_5’_ – are needed to fully specify the dinucleotide conformation. However, our results indicate that the positions of the local minima in the FES depends largely on the inter-base separation *R* and the dihedral twist angle *φ* and less on the tilt and roll angles. While all of the above structural parameters are specified in our calculations of the CD spectra and of structural and dynamical distribution functions, the visual representation of the FES is conveniently reported as a function of *R* and *φ*. The FES *G*(*R*, *φ*) of the dApdA dinucleotide shown in Fig. 2*A* applies to [NaCl] = 0.1 M, which is close to the monovalent salt concentration under physiological conditions. Using the same procedure, we also determined *G*(*R*, *φ*) at increasing salt concentrations (surfaces not shown). To test the validity of the FESs shown in Fig. 2*A*, we used the molecular configurations obtained from our 2 μs MD trajectories to calculate the CD spectra for dApdA. The results, as a function of salt concentration and using the procedures described in the Methods and the SI sections, are shown in Fig. 2*B*. Specifically, in the SI, Fig. S2 reports the angle that defines the direction of the electric dipole transition moment (EDTM) used in the CD calculations, while Table S1 reports the experimental values for the magnitudes and the molecular frame orientations of the EDTMs used in our calculations. In Fig. S3*A*, we show the favorable comparison between the CD spectra calculated using the point dipole approximation (PDA) versus the extended dipole model (EDM) for the Watson-Crick B-form conformation. We thus made use of the EDM for the majority of our CD calculations. In Fig. S3*B*, we show a comparison between the CD spectra calculated using spectroscopic parameters for the Adenine monomer obtained from different experimental studies. For all of our CD calculations, we made use of the experimental parameters obtained by Williams *et al*.,^7^ which we list in Table S2.

For the lowest salt concentration, [NaCl] = 0.1 M, we compared our calculations to the experimental CD spectrum of the dApdA dinucleotide obtained under these same conditions (see Fig. 2*B*).^15^ We obtained excellent agreement between the experimental and calculated CD in the long wavelength region of the spectrum (240 – 300 nm). We note that the agreement is less favorable in the short wavelength region (200 – 240 nm) of the spectrum, where the peak features are slightly blue-shifted and exhibit smaller amplitudes than the experiment. This latter disagreement is not surprising, given that the CD spectrum at shorter wavelengths is strongly perturbed by the high density of nearly degenerate electronic states, which makes the theoretical methods we employ in our calculations less accurate in this wavelength range.

In general, we find that the positions of the local minima within the free energy surfaces do not change with salt concentration, while their relative stabilities and equilibrium distributions do depend on this variable. The FES in Figs. 2*A* shows that the dApdA dinucleotide exists primarily as a mixture of the two chiral conformations with opposite handedness (*φ* = 40° and −80°) and nearly stacked inter-base separation *R* = 3.8 Å, together with an achiral conformation that shows no stacking of the bases (*φ* = 0°) and a significantly larger inter-base separation *R* = 4.7 Å. Henceforth, we will designate as ‘chiral’ a conformation that exhibits chiral stacking of the bases, and as ‘achiral’ conformations with no stacking of the bases, even though some components of the molecule, like the sugar, do of course retain their ‘chemical chirality.’

To study the effects of increasing salt concentration on the population of the chiral and achiral conformational states, we report in Fig. 2*C* the local probabilities calculated as the sum of the states contained within the areas of the FES defined by the red and white squares (panel *A*), respectively, for the chiral state with coordinates (3.8 Å, 40°) and for the achiral state with coordinates (4.7 Å, 0°) as a function of the salt concentration. We note that as the salt concentration is increased to [NaCl] = 0.5 M, the local probability of the chiral state with coordinates (3.8 Å, 40°) increases, while the weight of the achiral state slightly decreases. A further increase of the salt concentration to [NaCl] ~ 1 M begins to destabilize both of the stacked conformations in favor of the unstacked one, with the weight of the achiral state with coordinates (4.7 Å, 0°) strongly increasing. We observe a similar dependence on the salt concentration for the CD spectrum, which depends on the distribution of stacked bases. In Fig. 2*D*, we plot the difference CD spectrum for incremental changes of the salt concentration. For incremental increases of the salt concentration below [NaCl] = 1 M (0.1 → 0.5 M, 0.5 → 1.0 M), the difference CD spectrum shows little variation. However, for the incremental increase of 1.0 → 1.5 M, the difference CD spectrum undergoes a pronounced change. This change is also reflected by the value of the peak-to-peak amplitude of the difference CD spectrum (i.e., the difference between the positive peak value at 245 nm and the negative peak value at 270 nm), which is shown in the inset of Fig. 2*D*.

The above findings are in qualitative agreement with experiments involving the thermal melting of duplex DNA structures in NaCl, where increases in the concentration of monovalent ions tend to first stabilize the stacked conformation, resulting in an increase in the melting temperature. Then, at higher salt concentration (around [NaCl] = 1 M) this trend reverses, and the further addition of counterions slightly decreases the stability of the dsDNA conformation.^39–41^ For duplex DNA, this behavior is generally explained by assuming that an increase in salt concentration facilitates the screening of the negative charges situated on the phosphates in the DNA backbone, rendering the backbone more stable. However, at monovalent salt concentrations around 1 M, the concentration of ions in solutions becomes equivalent to the concentration of counterions closely bound to the phosphate backbone under ion condensation conditions. As a consequence, additional increases in salt concentration cannot further stabilize the double helix and other mechanisms (presumably ‘Hofmeister effects’^19,42,43,44,45^) come into play. Mechanisms involving the stabilization of long duplex DNA molecules by screening the repulsion between backbone phosphates cannot apply to dApdA, since only one phosphate is present. However, the counterions can alter the relative stabilities of the various conformations available to the dApdA dinucleotide by effectively neutralizing the negative charge of the single phosphate.

### Distributions of ions and water molecules around dinucleotides

To examine the roles of salt concentration on the observed structural transition we used the results of our MD simulations to calculate the distributions of the ions and water molecules of the solvent environment in the immediate vicinity of dApdA. This study provides physical insights into the origins of the changes in equilibrium base stacking conformations of this dinucleotide with increasing salt concentration. ^1–4,15^

The radial distribution function (RDF) of species *j* around species *i* is defined:

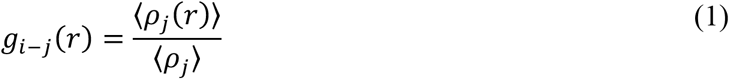

In Eq. (1), 〈*ρ*_*j*_〉 is the average density of species *j* and 〈*ρ*_*j*_(*r*)〉 is the local density of *j* within a spherical volume shell that is centered a distance *r* away from a molecule of species *i*. In what follows, we focus on the structure of the solvation shells as reflected by the RDFs between the phosphorous (P) atom of the negatively charged phosphate group of the dApdA molecule and the following solvent species: (i) the positively charged sodium ions (Na); (ii) the negatively charged chloride ions (Cl); and (iii) the hydrogen atoms of the water molecules (H).

Figs. 3*A* and 3*B* show the RDFs of the dApdA system at the lowest and highest salt concentrations we examined; i.e., [NaCl] = 0.1 and 1.5 M, respectively. The position-dependent oscillations of the RDFs reflect the local solvation shells of the water hydrogen atoms and of the ionic species relative to the central phosphate. At salt concentrations close to physiological conditions, ([NaCl] = 0.1 M Fig. 3*A*), the phosphate is coordinated with concentric ion shells, with the water hydrogen atoms forming interstitial layers between the shells. The RDF for water hydrogen atoms appears to be independent of salt concentration, with its first peak centered at *r* = 2.8 Å and its second peak at *r* = 4.2 Å. The RDFs for sodium and chloride ions, on the other hand, oscillate at half the spatial frequency of that of the water hydrogen atoms. The RDF for sodium ions has its first peak at *r* = 3.6 Å, which coincides with a trough for the water hydrogen atoms at this distance. Similarly, a trough for sodium ions occurs at *r* = 4.2 Å, which coincides with the second hydration shell for the water hydrogen atoms. The first ion shell for chloride ions occurs at *r* = 5.8 Å, which is the same position as the second ion shell for sodium ions. In general, the *n*th chloride ion shell occurs at approximately the same position as the (*n*+1)^th^ sodium ion shell, indicating that these ion shells have mixed compositions. As shown in Fig. 3*B*, the relatively well-defined boundaries between successive ion shells seen at the lowest salt concentrations become diffuse at the highest salt concentration tested ([NaCl] = 1.5M).

**Figure 3.**
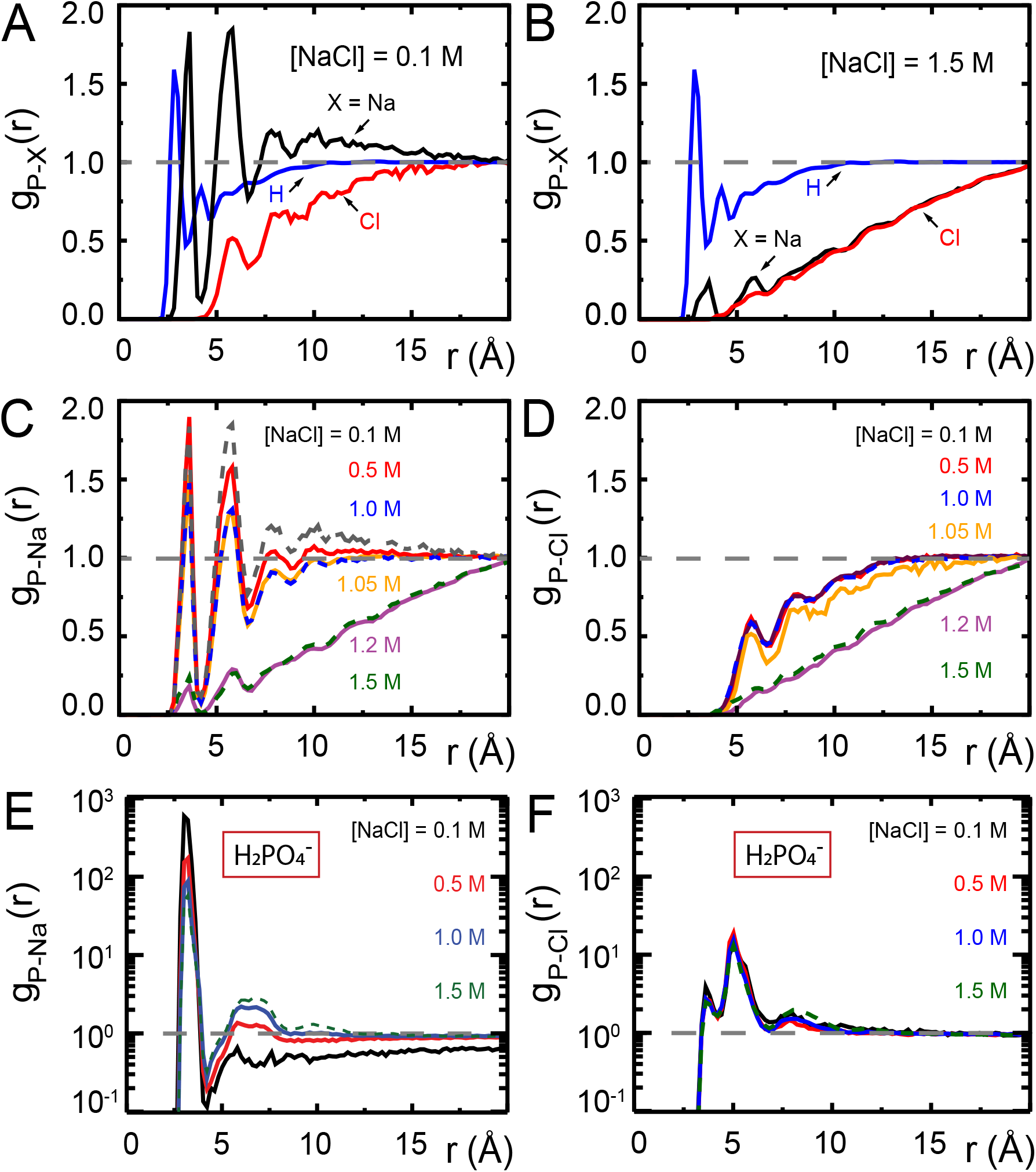
Radial distribution functions (RDFs) [Eq. (1)] obtained from MD simulations of dApdA between Na^+^, Cl^−^, and the H atoms of water and the P atom of the anionic phosphate of the dApdA dinucleotide at salt concentrations ([NaCl]) of (***A***) 0.1 M and (***B***) 1.5 M. RDFs for (***C***) sodium ions and (***D***) chloride ions over the range of salt concentrations [NaCl] = 0.1, 0.5, 1.0, 1.05, 1.2, and 1.5 M. RDFs for sodium (**E**) and chloride (**F**) ions obtained from MD simulations of the phosphate anion 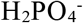 at the salt concentrations ([NaCl = 0.1 (black), 0.5 (red), 1.0 (blue), 1.5 M (light blue). Unlike the RDF plots for dApdA, the RDFs of 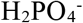 in aqueous solutions do not show the sharp change in the ion shell structure at [NaCl] ≅ 1. 0 M (see also text).

Our observation of a well-ordered structure of successive ion shells at low salt concentration is largely consistent with simple models of counterion condensation, which is an important contributing factor to the stability of larger nucleic acid molecules.^46,47^ Figs. 3*C* and 3*D* show, respectively, the RDFs of sodium and chloride ions, each as a function of salt concentration. For both ions, the RDFs appear to change little over salt concentrations between [NaCl] = 0.1 – 1.0 M, yet exhibit an abrupt loss of ion shell structure at salt concentrations slightly greater than 1.0 M.

To illuminate the role(s) of the adenine bases in this situation, we performed a set of 400 ns simulations of 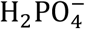 at increasing monovalent salt concentration ([NaCl]=0.1, 0.5, 1.0 and 1.5 M), and studied the ion distributions around a singly-charged phosphate ion, 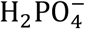, in aqueous solution (see Figs. 3*E* and 3*F*). In 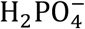 we observed an alternating structure of positive and negative ion shells consistent with simple models of counterion condensation. However, we found no signature of the abrupt disruption of the ion shell structure at salt concentrations greater than 1.0 M that was observed with the dApdA dinucleotide.

We next turned our attention to a closer examination of the solvent orientation around dApdA. As mentioned previously, the structure of the water, which is reported as the position-dependent RDF of the water hydrogen atoms relative to P, *g*_P-H_(*r*), does not change significantly with salt concentration (see Figs. 3*A* and 3*B*). More detailed behavior is observed in the position-dependent orientational distribution function (ODF) of the water dipole moment as a function of its separation from the central P atom. The ODF is defined as the average cosine, 〈cos *θ*(*r*)〉, of the angle *θ* that subtends the permanent dipole moment of the water molecule and the vector connecting the P atom to the water O atom, 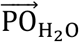, as shown in Fig. 4*A*.

**Figure 4.**
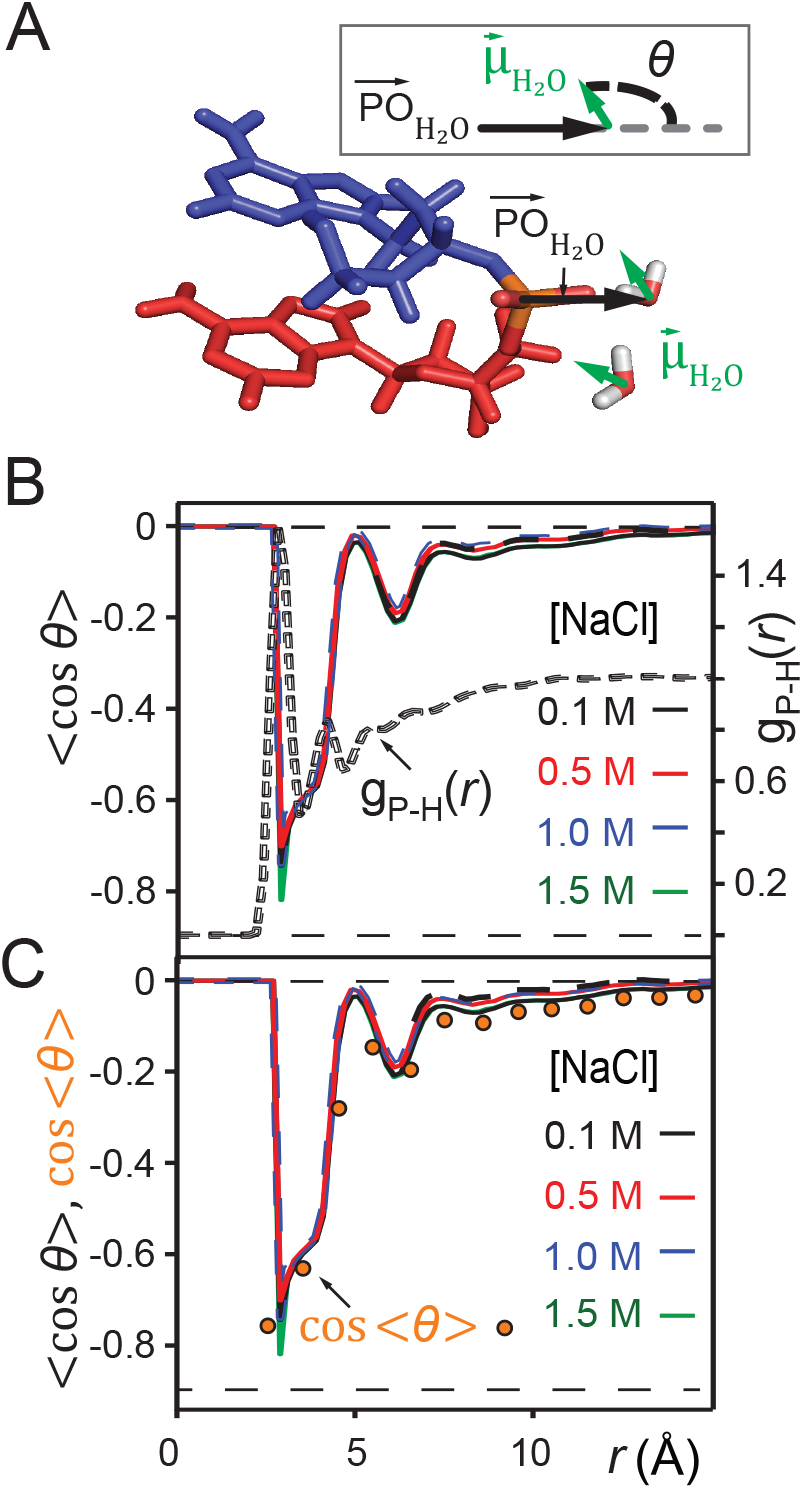
*(A)* Definition of the angle ***θ***, which subtends the permanent dipole moment of the water molecule 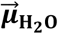 and the vector 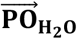 connecting the phosphorous atom to the oxygen atom of the water molecule. (*B*) Orientation distribution functions (ODFs) for the dipole of the water molecule relative to the phosphate-oxygen (water) bond and RDF of the hydrogen of water, ***g***(***r***)_***P-H***_ >, of Fig. 3*A*. *(C)* Superimposed on the ODFs defined as < ***cos θ*** >, are the orange points indicating the cosine of the average angle, ***cos*** < ***θ*** >.

Fig. 4*B* shows the ODFs of water relative to the central phosphate of dApdA as a function of salt concentration. It also shows the RDF of the water hydrogen atoms. The position dependence of the ODFs shown in Fig. 4*B* exhibits damped oscillations that vary across successive hydration layers for all salt concentrations, similar to the behavior observed for the ion shell structures shown in Fig. 3. The ODFs show a sharply pronounced feature centered at *r* = 2.8 Å, which is coincident with the first peak of the RDF. The shapes of the underlying distributions of the angle *θ* within a narrow range of distances *r* ensures that 〈cos *θ*〉 ≈ cos〈*θ*〉 (orange points in Fig. 4B). The distributions of the angle *θ* for a given hydration shell, with each distribution corresponding to one orange point in Fig. 4B, are reported in Fig. S8 of the SI. Thus, the narrow feature at *r* = 2.8 Å has an approximate peak value of cos〈*θ*〉 = –0.8, which that the water H atoms within this first indicates hydration shell are highly oriented with dipole moment 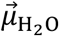 directed toward the central P. Furthermore, the presence of the broadened ^0^ shoulder centered near the second hydration layer (at *r* = 4.2 Å), with peak value approximately cos〈*θ*〉 = –0.6, indicates the preferential orientation the O-H bond vectors of water molecules within the second hydration shell towards the oxygens of water molecules within the first hydration shell. between water molecules of the first and second hydration shells are stronger than the Coulomb interaction between the negatively charged phosphate and the water dipole moments of the We thus see that hydrogen bonding interactions second hydration shell. We further note that the distribution of angles *θ* over a given range of distances *r* broadens nonuniformly as the distance from the central P increases, indicating the presence of hydrogen bonding between successive hydration layers and the ensuing loss of orientational correlation between the water dipoles and the central P. At the separation *r* = 5 Å, the values of the ODFs are approximately zero, indicating the absence of orientational alignment. A recurrence of partial orientational order occurs at separation *r* = 6 Å, which appears to coincide approximately with the position of the first ion shell of the Cl-ions.

We note that the ODF exhibits a weak, but clear, dependence on the salt concentration. For the case of [NaCl] = 0.1 M, the sharp feature at *r* = 2.8 Å indicates a pronounced orientation, which becomes slightly less ordered for [NaCl] = 0.5 M. At the higher salt concentrations of [NaCl] = 1.0 M and 1.5 M, the orientation of the water dipole moments become slightly more ordered. The changes in the ODF of the water molecules as a function of increasing ion concentration are small. Rather, the leading factor in determining the stabilities of the conformations of the dinucleotide structure in solution appears to involve the distribution of monovalent ions, and the modification of this distribution with increasing ion concentrations (see Figs. *3A – D*). Water, however, does appear to play a role through its orientation, which is both distance and weakly salt-concentration-dependent. Interestingly, this study also shows that the stabilization of dApdA stacking by increasing counterion concentrations, and the observed sharp transition of the ion structure around 1 M are dependent on the presence of the bases of the dApdA dinucleotide, and do not occur when the ionized phosphate molecule is present alone (Figs. *3E – F*). The consistency of the observed trend with the effects of increasing salt concentration on the experimental melting curves of DNA suggests that this ion-related base stacking mechanism of DNA stabilization is already present and operational, even at the level of the isolated dinucleotide.

### Markov state model analysis of the free energy landscapes of the dApdA dinucleotide and comparison with CD spectral analysis

The theoretical representation of the CD spectrum for a flexible molecule in solution is the summation of contributions from the myriad microscopic conformational states (i.e. microstates) that exist at equilibrium. Intuitively, we expect the dApdA dinucleotide to fluctuate between various ‘open’ and ‘closed’ base conformations, which in turn are stabilized (or destabilized) by the surrounding hydration and ion shells. We first determined the CD spectrum by summing over equally weighted contributions from the 10 million microstates that are sampled from each of our MD simulations (see Fig. 2*B*). Although the above ‘brute-force’ approach is straightforward, it suffers from two significant limitations: (i) it provides little insight into the interpretation of CD in terms of specific molecular conformations; and (ii) it becomes computationally inefficient if one adopts more sophisticated quantum chemical models to calculate the CD spectrum beyond the extended dipole model used here, because one would need to perform advanced calculations for each of the 10 million microstates. In reality, only a relatively small subset of the total number of possible conformational states is expected to contribute significantly to the measured CD spectrum. The specific states that dominate the CD are the stacked and chiral conformations of the dinucleotide, for which both the electronic coupling between monomer electric dipole transition moments (or EDTMs) and the rotational strengths resulting from these couplings are significant (see SI). Conformational states that are unstacked, in addition to those that are stacked and essentially achiral, contribute much less to the CD spectrum To determine the dApdA configurations that are most relevant for the interpretation of the CD spectra, we used a Markov state model (MSM) analysis^20–22^ to subdivide the 10 million microstates obtained from our MD simulations into a relatively small number (five) of ‘macrostates’, each of which is associated with a distinctive region of the free energy landscape (see Fig. 5*A*). Each macrostate represents a collection of conformationally related states, or ‘microstates’, which rapidly interconvert during the simulation, while slow transitions between macrostates require crossing large energy barriers. Starting from the MSM analysis, we calculated the CD spectrum as the sum of those, unequally weighted, macrostates.

**Figure 5.**
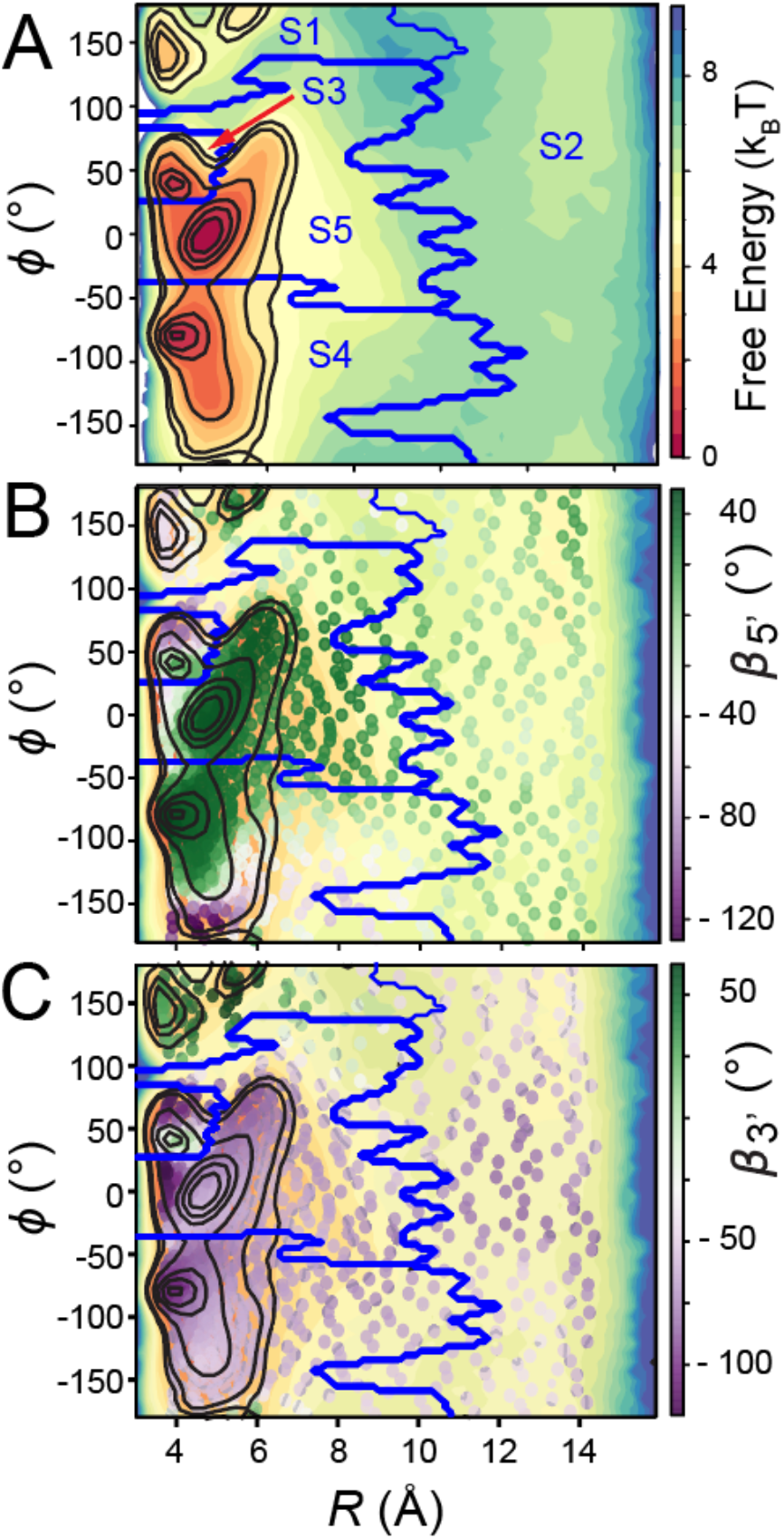
(***A***) The free energy landscape *G*(*R*, *φ*) of the dApdA dinucleotide (shown in Fig. 2*A*) is sub-divided by dark blue boundaries into five regions (labeled S1 – S5), which are called ‘macrostates.’ The macrostate assignments were derived by performing a Markov state model (MSM) analysis of MD simulation data for [NaCl] = 0.1 M. The *anti* (Watson-Crick) form conformation is contained within the boundaries of the S3 macrostate, while the *syn* (Hoogsteen) containing form is included within the boundaries of the S4 macrostate. Superimposed on the free energy landscape *G*(*R*, *φ*) of the dApdA dinucleotide we show the orientation of the 5’ base (***B***) and of the 3’ base (***C***), respectively. The macrostate S3, which contains the *anti* form conformation, correctly displays both bases with positive orientation (green free energy minimum of S3 in both panels *B* and *C*), while macrostates S4 and S5, which contain a *syn* base presents the 3’ base flipped with respect to the 5’ base (green in panel *B* and purple in panel *C*).

Thus, the kinetic processes that occur in the simulations are partitioned between those that occur faster than the ‘lag time’ (in this study τ = 500 ps) and those that occur more slowly than this time scale. Transitions between conformations within a given macrostate occur frequently and are non-Markovian, while transitions between conformations belonging to different macrostates occur less frequently and are Markovian. The ‘lag time’ is defined as the time that fulfills the above-stated condition of Markovian transitions between conformations belonging to different macrostates (for details see SI and, in particular, Fig. S6, which tests the Chapman-Kolmogorov condition, thus ensuring the Markovian nature of our partitioning of the FES into five states. Table S3 shows that the MSM analysis is insensitive to the choice of the number of microstates).

To further reduce the computational requirements for the calculation of the CD spectrum, we identified one averaged structure, together with its relative weight, for each of the five key macrostates that are relevant to the CD observable. We found that the total CD spectrum can be accurately represented by the weighted sum of the contributions from these five averaged structures (see SI), which could be used for modeling of the CD spectrum using more advanced quantum chemical models.

In Fig. 5*A*, we show the free energy landscape for the dApdA dinucleotide monophosphate in 0.1 M salt (NaCl), and its subdivision into five macrostates (indicated by dark blue contour lines and labeled S1 – S5), which we established using our MSM analysis approach. Each of the five macrostates exhibits qualitatively different behavior in terms of the relative stabilities of the dinucleotide conformation. The S1 macrostate includes 264,608 microstate configurations (2.6% of the total 10 million) and is dominated by a relatively shallow free energy basin with a narrow range of values for the inter-base separation *R* < 6 Å and relative twist angle: 100° < *φ* < 180°. The S2 macrostate, on the other hand, describes a relatively broad and featureless region of the free energy landscape, which encompasses a wide range of ‘open’ and ‘unstacked’ values for the inter-base separation (*R* >10 Å) and unrestricted twist angle: −180° < *φ* < 180°. Like the S1 macrostate, the S2 macrostate represents a minority of the total population, with just 276,786 microstates (2.8% of the total 10 million). The majority of the total conformation population is contained in the combined S3, S4 and S5 macrostates, with the number of microstates in S3: 895,636 (9.0%); in S4: 3,729,206 (37.3%); and in S5: 4,833,758 (48.3%). Moreover, the S3 macrostate contains the free energy minimum with *R* = 3.8 Å and *φ* = 40°, the S4 macrostate contains a minimum with *R* = 3.8 Å and *φ* = −80°, and the S5 macrostate contains a minimum with *R* = 4.7 Å and *φ* = 0°. We thus identify the S3 macrostate with an ensemble of stacked right-handed chiral conformations that include the *anti* form; the S4 macrostate with an ensemble of stacked left-handed chiral conformations, including the precursor of the Hoogsteen structure; and the S5 macrostate with one slightly less stacked and more achiral conformation, which also includes a *syn* structure.^30–37^ The borders between macrostates show a ‘fine structure’ that represents the maximum of the energy at the top of the free energy barriers, where the states are less frequently sampled by the simulation. Thus, the high energy regions in the free energy map may display roughness, which can be smoothed to avoid overfitting.^48,49^ However this step is not needed in our study because the results of our analysis depend largely on the minima of the free energy maps, which are statistically well sampled.

To confirm the presence of Hoogsteen-like structures in the S4 and S5 regions, we present – in Figs. 5*B* and 5*C* – a study of the roll angles *β*_5’_ and *β*_3’_, for the 5’ and 3’ base, respectively. It is known, for structures containing a Hoogsteen conformation, that one of the two bases in the dApdA dinucleotide is ‘flipped’ relative to the ‘standard’ conformation characteristic of the Watson-Crick geometry. Figure 5*B* shows that in the S3 macrostate the most stable structures have a positive roll angle *β*_5’_ for the 5’ base (green). Figure 5*C* shows, instead, that while the roll angle for the 3’ base is still positive in the microstate S3, the same 3’ base is flipped in microstates S4 and S5 (purple), confirming the presence of a *syn* Hoogsteen-like conformation in macrostates S4 and S5, and the *anti* Watson-Crick-like form in macrostate S3.

In Figs. 6*A – E*, we compare the experimental CD spectrum (dashed black curve) to our CD calculations corresponding to each of the five macrostates (blue curves), which are based on summing over the microstate configurations that lie within the partitioned boundaries of the free energy landscape shown in Fig. 5*A*. We also show the proportionately weighted contribution of each macrostate to the CD spectrum (red). From Figs. 6*A* and 6*B*, we see that the S1 and S2 macrostates, which represent minority fractions of the total population (2.6% and 2.8%, respectively), do not contribute significantly to the CD spectrum.

**Figure 6.**
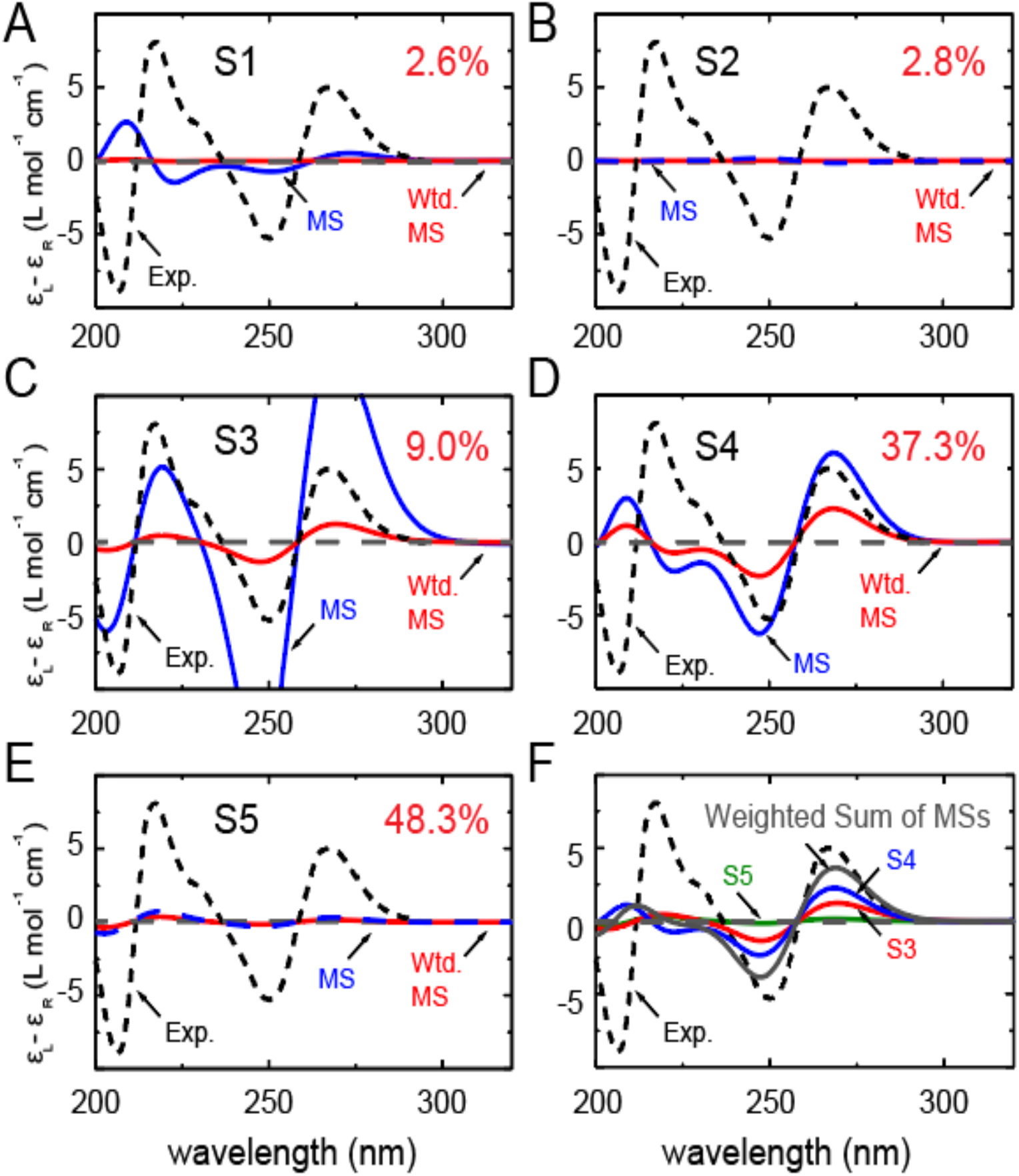
Macrostate decomposition of the CD spectrum of the dApdA dinucleotide by Markov state model (MSM) analysis of MD simulation data for [NaCl] = 0.1 M. The total CD spectrum is calculated from 10 million MD frames (or microstates), and the component spectra for macrostates (***A***) S1, (***B***) S2, (***C***) S3, (***D***) S4 and (***E***) S5 constitute 2.6, 2.8, 9.0, 37.3 and 48.3% of the total CD spectrum, respectively. The component CD spectra for each macrostate are shown in blue, and the number-fraction weighted contributions are shown in red. (***F***) The sum of number-fraction weighted macrostate contributions to the total CD is shown in gray. Also shown separately are the number-weighted contributions of macrostates S3, S4 and S5 (green, blue and red, respectively). In all panels, the experimental CD spectrum (from (12)) is shown as dashed black curves.

Similarly, the S3 macrostate (Fig. 6*C*), which contains the coordinates of the B-form conformation, also represents a comparably small fraction of the total population (9.0%). On the other hand, the S4 macrostate (Fig. 6*D*) contains a significant fraction of the total population (37.3%) and is largely composed of left-handed base-stacked conformations, which gives rise to a strong CD signal. We note that the calculated CD spectrum of the S4 macrostate has a similar ‘right-handed’ shape (in the long wavelength regime) to that of the S3 macrostate, in spite of their apparent opposite chiral symmetries. This is consistent with the *syn* structure (i.e., with roll angles *β*_3’_ ≈ 180° and *β*_5’_ ≈ 0°). The detailed calculation of the spectra for all five macrostates is reported in Fig. S4 of the SI, which shows the spectral decomposition of the degenerate CD spectrum for the average structure of each macrostate. From the spectral decomposition it is straightforward to see that the flipping of one base is responsible for a CD spectrum that is consistent in the Watson-Crick structure and in the *syn* conformation of dApdA. We note that this behavior may not be observed in dinucleotides with different base compositions, because the transition dipoles are different. Although the S5 macrostate (Fig. 6*E*) represents the highest fraction of the total population (48.3%), it is dominated by an achiral and slightly unstacked *syn* conformation which, because of its symmetry, results in a negligible CD contribution to the total spectrum. In Fig. 6*F*, we show the individual weighted contributions for the S3 (9.0%), S4 (37.3%) and S5 (48.3%) macrostates, in addition to the weighted sum of all the macrostates (gray curve). We thus see that the favorable agreement we observe between experiment and theory in the long wavelength regime is essentially the result of two significant contributions, a minor contribution from macrostate S3 and a larger contribution from macrostate S4.

Having identified the key macrostates relevant to the CD observable, we used this information to determine the smallest number of structural parameters necessary to characterize these macrostates. We thus identified five averaged conformations, one for each macrostate, which, properly weighted, were used to calculate the CD spectrum. The comparison between the contribution to the CD spectrum from all the conformational states in a macrostate and the contribution from the averaged macrostate structure are shown in Fig. S6 of the SI, with structural parameters listed in Table S6. The calculation of the CD spectrum with only five conformations is in good agreement with the complete calculation, while it greatly speeds up the computation time needed to calculate the CD spectrum. In principle, such structural models can be used for the general interpretation of any spectroscopic measurement performed on the dApdA system.

In reconsidering the previous interpretations of the CD spectrum by Lowe and Schellman, and given that the signal from the un-stacked mono-nucleotide is comparatively negligible, our study suggests that the stacked ‘native’ form of the dinucleotide is primarily given by the sum of the S3 and S4 states, because the S1 state is less densely populated. Analogously, the unstacked ‘denaturate’ state corresponds in this study to the S5 state, which is more populated than the unstacked S2 state.

The large degree of conformational disorder that characterizes macrostate S2 contrasts with the highly ordered macrostates S3 and S4. The stabilities of macrostates S3 and S4, relative to macrostate S5, are reminiscent of the ‘solvophobic’ models for nucleic acid base stacking,^2,11,12^ in which the ‘stacked’ macrostates S3 and S4 are favored due to enthalpic base stacking interactions, ^50^ which offsets the configurational entropy of the disordered S2 macrostate. Solvophobic base stacking is known to be favored by a decrease in the enthalpy Δ*H* and opposed by a decrease in the entropy Δ*S*. Solvophobic bonding, as defined here, is ‘enthalpically driven’ and differs significantly from hydrophobic bonding, which is generally thought to drive protein folding^51^ by a positive change in the entropy of the system. Such physical models are supported by studies that examine the stabilizing and destabilizing effects on base stacking by various salts and other solvent additives.^10,18^

### Mean first passage times (MFPTs) for dApdA macrostates and pathways of macrostate interconversion

While CD spectra provide a useful measure of the stationary (equilibrium) properties of the dApdA system, they do not provide information about the dynamic processes involved in state-to-state interconversion. In this section we apply the results of our MSM analysis of MD trajectories to the investigation of the kinetic pathways associated with the free energy landscape, and to identify pathways of interconversion between the various stacked and unstacked macrostates.

To characterize the kinetic properties of the dApdA system, we examined the mean first passage times (MFPTs) of the five macrostates, which are assigned to the regions of the free energy landscape shown in Fig. 5*A*. The MFPT τ_*i*→*f*_ is the average time for the system to undergo a transition to state *f*, provided that it was initially in state *i*.^21,52^ We determined the MFPTs for the free energy landscape of dApdA at salt concentration [NaCl] = 0.1 M (see full data set in Table S4 of the SI. Also, Table S5 shows that the MFPTs are insensitive to the number of microstates selected in the MSM analysis). Macrostate S2 represents the region of the free energy landscape with the greatest degree of conformational disorder; thus, it can be considered to serve as an end-state for base-unstacking.

Moreover, while macrostate S3 is approximately B-form in character, the relative roll angles of macrostates S4 and S5 are greater than 90°, which in each case corresponds to a base configuration that has been flipped into the Hoogsteen-like conformation. Thus, the process of ‘base-flipping’ may play an important role in the dynamics of the dApdA system, although in longer strands of (especially) duplex DNA, such flipping may be suppressed by the overall cooperativity that controls the order-disorder transitions for these larger macromolecular species. Nevertheless, these studies of the less cooperatively stabilized dinucleotide may provide insight into structural rearrangements that in principle could, and likely – with some frequency – do, occur in larger biologically relevant DNA macromolecules.

We used the transition path sampling (TPS) method^25–29^ to determine the frequency of events in which an initially base-stacked macrostate (e.g., S3 or S4) undergoes successive conformational changes that permit entry into the region of the free energy landscape characterized by the ‘final’ unstacked macrostate S2. When the system initially occupies macrostate S3, which corresponds to the average base stacking of the Watson-Crick B-form, we found that the dominant pathway leading to macrostate S2 (with 46% probability) was S3 → S5 → S2. Thus, base-unstacking from the right-handed B-form conformation occurs predominantly by a two-step process through the achiral S5 intermediate, in which one of the adenine bases has been flipped. The remaining, less prevalent base-unstacking pathways were S3 → S4 → S2 (with 26% probability); S3 → S5 → S4 → S2 (with 15% probability); and S3 → S2 (with 10% probability). When the system occupied initially the left-handed and base-flipped macrostate S4, which corresponds to a Hoogsteen base-stacking configuration, the two most prevalent unstacking pathways were the one-step S4 → S2 pathway (with 47% probability) and the two-step S4 → S5 → S2 pathway (with 40% probability).

We see that, in general, transitions to the most sparsely populated macrostates S1 and S2 occur relatively slowly (in ~35 to 60 ns), while transitions to the most highly populated macrostate S5 occur relatively quickly (in ~2 to 5 ns), suggesting that the macrostate S5 acts as a common intermediate for the pathways between the other macrostates for the stacking-unstacking transition.

Because the energy barriers in dApdA are small, the height(s) of the barrier(s) that the system has to overcome to transition between any two macrostates is close to the difference in free energy between the two states. Thus, the kinetics of the interconversion between macrostates are driven primarily by their relative stabilities. It is reasonable to expect that cooperativity in base stacking will increase the heights of the energy barriers between conformational states in both the ssDNA and the dsDNA. Such information can provide new insights into the mechanisms of base stacking-unstacking transitions in nucleic acids and the possible role of these processes in biologically important protein-nucleic acid interactions.

## Discussion

### Structural and dynamic characterization of ‘breathing’ fluctuations at the dinucleotide level

Thermally activated breathing fluctuations, in which flanking nucleic acid bases spontaneously move away from their stacked and hydrogen-bonded conformations, are thought to be important initial steps in the pathways that lead to DNA denaturation and the specific binding of proteins to DNA.^1–6^ Despite their relevance, the details of the interactions and kinetics that control breathing fluctuations are still largely not understood. It is known, however, that the stacking interactions of the bases within nucleic acids are the dominant stabilizing forces of the native conformations that oppose the melting of DNA, while inter-strand base-base hydrogen bonding and cooperativity play less important stabilizing roles.^10,17,50^ Traditionally the equilibrium between stacked and unstacked base conformations has been studied by circular dichroism (CD) experiments, which are sensitive to the conformational chirality of the base stacking.^15^ Such measurements, however, are limited in the amount of information that they can provide because CD spectra cannot be directly inverted to determine the conformations that contribute to these spectroscopic signals.

CD spectroscopy is an important biophysical tool for the analysis of nucleic acid structure, in that the relationship between CD spectra and local nucleic acid base conformation can be understood in terms of quantum chemical principles. Nevertheless, for many of these systems the free energy landscape can favor the simultaneous presence of multiple conformations at equilibrium, many of which may interconvert due to thermal fluctuations. Thus, the complexity of the free energy landscapes of nucleic acid systems is a significant obstacle for achieving a meaningful interpretation of CD spectra.

### Solvophobic effects on the conformational stability of dinucleotides

Early studies by Lowe and Schellman of the base stacking-unstacking equilibrium focused on the CD spectrum of the dApdA dinucleotide monophosphate as a function of increasing monovalent salt concentration, because the stacking interactions of the elementary dinucleotide unit could be isolated and studied independently of other stabilizing factors.^10^ These studies concluded that the stacking-unstacking equilibrium of dinucleotides can be modeled as a two state transition, where the driving force for the stacking of the bases is ‘solvophobic’ in nature; i.e., driven by a decrease in the enthalpy of the process (Δ*H* ≈ −6.6 kcal mol^-1^ at *T* = 293K), and opposed by a decrease in the entropy of the system (Δ*S* = −23 e.u. such that −*T*Δ*S* ≈ 6.7 kcal mol^-1^ at *T* = 293K).^10,14,15,50^ Thus, these workers concluded that the transition as a whole was likely driven – to a major extent – by rearrangements of the molecules of the solvent environment present (here water molecules and ionic species). However, these studies could not exclude the possibility that more than two states might contribute to the overall CD signal, and thus could not define the precise nature of the underlying conformations.^10^ They did determine, however, that each of the two states of the dApdA dinucleotide that contributed to the CD signal was most likely present as a number of similar configurations, and that the state with highest disorder and entropy was likely to be more stable at high temperatures and at higher monovalent salt concentrations.

### Conclusions and overview

In the present study we have established a methodology that can be used to relate the CD spectrum to the underlying relevant molecular conformations. We combined extensive MD simulations (μs in duration) with direct calculations of the CD spectrum. Our CD calculations were based on standard methods^24^ and an extended-dipole model (EDM)^53^ to estimate the exciton coupling between the electric dipole transition moments (EDTMs) of the adenine bases of dApdA. The EDM takes into account the finite length of the electronic transition charge distribution across the adenine chromophore, and it correctly describes the dependence of the electronic coupling on the inter-base twist angle *φ* and the relative tilt angle *β*_*S’*_ − *β*_*S’*_ (coordinates defined in Fig. 1). By calculating the CD spectrum for each of the 10 million conformations in the MD simulations, we obtained excellent agreement between our CD calculations and previously published experimental spectra of the dApdA system at approximately physiological salt concentration [NaCl] = 0.1 M.^10,15^ Nevertheless, the calculation of the CD spectrum by these procedures provided little insight into the important conformational states contributing to the CD spectrum, and can become computationally too expensive if sophisticated quantum chemical calculations are adopted to calculate the exciton couplings from the detailed electronic structure of the adenine bases.

To surmount this problem, we performed a Markov State Modeling (MSM) analysis of the free energy landscape of the dApdA dinucleotide and identified five kinetically separable macrostates, each containing conformational species that can rapidly interconvert. We then calculated a single averaged conformation to represent each of the five MSM macrostates, and we found that the total CD spectrum can be represented accurately by the weighted sum of the contributions from the averaged structures of these macrostates.

We found that only two states exhibit both stacked and chiral conformations, which are necessary to provide significant exciton coupling between monomer EDTMs and rotational strengths, thus contributing to the CD observable. The two states are conformational ensembles with opposite chirality, which contain the *anti* (Watson-Crick B) form (S3) and a *syn* (Hoogsteen) flipped-base conformation (S4), respectively. A third highly populated state is an achiral *syn* state, with a slightly unstacked conformation (S5) that does not contribute significantly to the CD signal. We observed that both the S3 and the S4 states provide right-handed CD features in the long wavelength region of the dApdA spectrum. These results are qualitatively consistent with the early hypothesis that two leading states dominate the CD spectrum, but now provide more detailed information about the nature of those states.^10^ We conclude that both the S3 and the S4 states contribute to the stacked conformation detected by Lowe and Schellman, while their unstacked conformation likely comprises the S5 state, which is the most populated, and to a lesser extent the fully unstacked state S2. Furthermore, our study shows that the Hoogsteen structure plays a key role in the mechanism of the stacking and unstacking pathways of the bases in dApdA, and possibly in DNA as well, as it is present in the highly populated stacked S4 and unstacked S5 conformations at all salt concentrations.

By connecting the CD spectrum of the dApdA dinucleotide to five leading conformational states as a function of salt concentration, we were able to obtain information about how the distribution of ion shell structures affects local base-stacking interactions. In agreement with early experiments, we observed that the effect of increasing salt was to decrease the magnitude of the CD signal over the 240 – 300 nm regime, and that the decrease in CD signal is accompanied by a shift in the equilibrium population of open (unstacked) achiral conformations relative to closed chiral conformations. Our findings show an initial increase of the base stacking stability with increasing monovalent salt concentration (NaCl or KCl), followed by a decrease of stability at salt concentrations higher than 1.0 M. By analyzing radial distribution functions and orientation distribution functions of the ions and water solvent, respectively, we observed that the changes in local base stacking conformation at high salt concentrations are correlated strongly with the disruption of the ion shell boundaries, and weakly with a change in the orientations of the water dipole moments. Over the full range of salt concentrations, the orientations of water molecules within successive hydration shells are highly correlated, from layer to layer, through hydrogen bonding. Thus, the relatively large negative change in solvent entropy is attributed to the emergence of order of the ion shell structure upon base stacking, rather than the restructuring of the water dipole moments. These findings provide a more detailed picture in the context of the solvophobic bonding model, in which the enthalpic base stacking interaction is closely balanced by the decrease in entropy of the solvent environment. In contrast, this behavior is not observed for the singly-charged phosphate anion in isolation, suggesting that the presence of the bases in dApdA structure may be responsible for the disruption of the ion shell structure upon base unstacking.

We also find that the trend in base stacking stability with increasing monovalent ion concentration for the dApdA dinucleotide is consistent with trends observed for the more complex duplex DNA.^16,39^ Although other factors, such as H-bonding and the cooperative stacking of multiple bases are known to play an important role in determining the stability of dsDNA structures, our results suggest that the restructuring of the ion shells about the central phosphate ion with increasing salt concentration, observed in dApdA, may also play a role in regulating the stability of larger DNA macromolecules.

## Data availability

Data discussed in the paper have been made available in Zenodo with reference number 3971255.

Source codes for the calculations of the CD spectra using the Extended Dipole Approximation and the Point Dipole Approximation have been made available in GitHub at https://github.com/GuenzaLab/dApdA.

## Acknowledgements

The authors are grateful to Dr. Pablo G. Romano for his contributions in early phases of this project. We are also grateful for many helpful discussions with the members of the Guenza, Marcus and von Hippel research groups.

## Supporting Information: Dinucleotides as simple models of the base stacking-unstacking component of DNA ‘breathing’ mechanisms

### I. MOLECULAR DYNAMICS SIMULATIONS OF THE dApdA DINUCLEOTIDE

Molecular dynamics (MD) simulations of the dApdA dinucleotide monophosphate molecule in aqueous solution were performed at increasing salt concentrations ([NaCl]= 0.1, 0.5, 1.0, 1.05, 1.2 and 1.5 M) in the NPT ensemble using the GROMACS software program.^1^ The length of the simulation box was allowed to fluctuate, so that the average distance between the box boundary and the dApdA molecule was approximately 20 Å. The initial configuration for the dApdA dinucleotide was selected as the B-form conformation, for which we obtained atomic coordinates from the ambertools software package [http://casegroup.rutgers.edu/].

**Figure S1.**
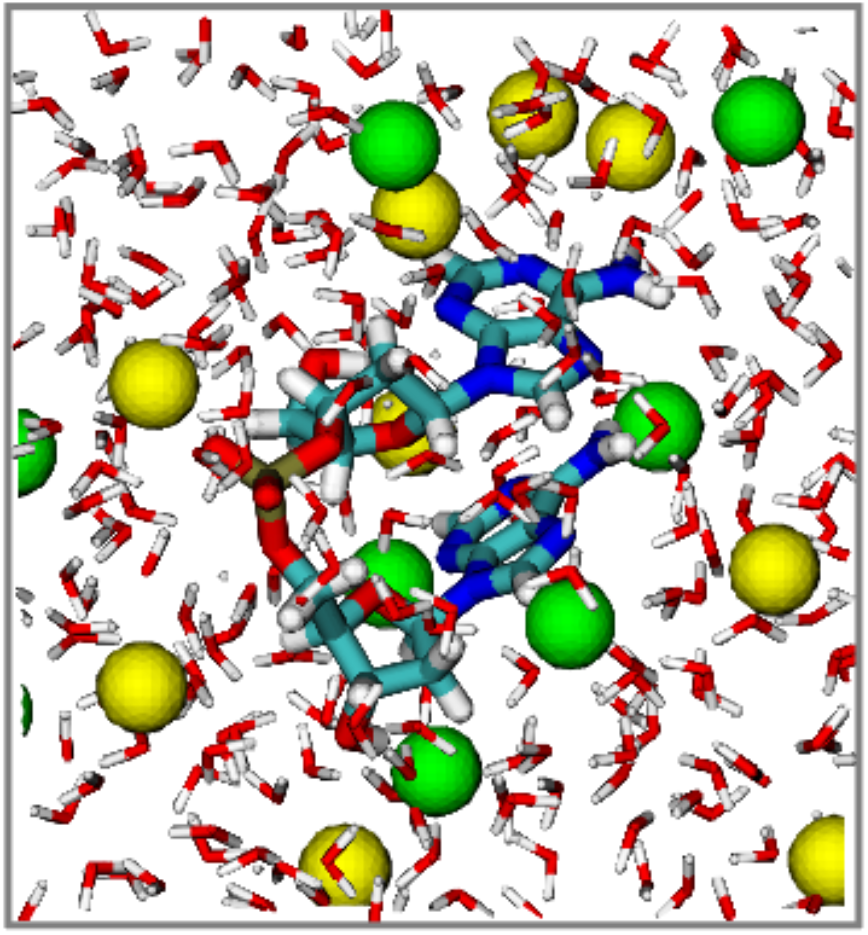
A sample configuration frame taken from an MD simulation run of dApdA dinucleotide monophosphate in TIP3P water with [NaCl] = 0.1 M. Sodium ions are shown as yellow spheres, and chloride ions as green spheres. The atoms of the dApdA and water are colored according to CPK rules, except for carbon, which is colored light blue.

Simulations were performed with the Amber03 force-field^2^ and the TIP3P water model^3^ to model the dApdA molecule and the water component of the solvent, respectively. While these models were not specifically parameterized to achieve accurate CD calculations of the dApdA dinucleotide, they have been used successfully for nucleic acid systems in the past and represent the state-of-the-art for simulations of DNA in solution. A sufficient number of sodium and chloride ions were included to achieve the target salt concentration. The energy of the solvated structure was minimized using the Steepest Descent algorithm for 500 steps. The system was then heated to 300 K and equilibrated in the isothermal-isobaric (NPT) ensemble using a time step of 2 fs over a period between 50 − 100 ns.

Production runs at each salt concentration were performed for a total duration of 2 μs in the NPT ensemble in order to ensure sufficient sampling of the conformational landscape. These simulations used the stochastic velocity-rescaling thermostat^4^ with a time constant of 0.2 ps, and the Parrinello-Rahman barostat (using an isotropic pressure coupling time constant of 1.0 ps). We implemented the Leap-Frog algorithm to integrate Newton’s equations of motion using the LINCS constraints fourth order in the expansion of the constraint coupling matrix, which included one iteration to correct for rotational lengthening.^5^ We set the time step to 2 fs, and truncated the Lennard-Jones interactions using a cutoff distance of 10.0 Å. We additionally used a particle mesh Ewald sum to handle long-range electrostatic interactions with a real space cutoff of 10.0 Å and a grid spacing of 1.0 Å. The Verlet neighbor list algorithm was applied with a frequency of 10 MD steps to enhance the computational speed. Trajectory frames were stored every 0.2 ps. In Fig. S1, we show a sample frame from one of our MD trajectories. At each salt concentration we included ~ 10 million such frames in our CD calculations for the dApdA system.

### II. CALCULATIONS OF CIRCULAR DICHROISM (CD) SPECTRA FROM MOLECULAR CONFIGURATIONS

We applied the standard methods developed by Schellman and others to model the delocalized electronic states of the dApdA dinucleotide as a function of base stacking conformations.^6,7,8^ In the formalism that follows, we consider only the contribution to the CD spectrum that emerges from the exciton interactions between the component adenine bases of the dApdA dinucleotide, and we neglect the minor contribution to the CD from the non-interacting adenine monomer, which provides a relatively weak signal for the fully unstacked conformation. When light of frequency *v* interacts with a solution of optically active molecular chromophores, the left and right circularly polarized components are absorbed to different extents. The frequency (or wavelength) dependence of the differential extinction between left and right circular polarizations, Δ*ε*(*v*) = *ε*_*L*_(*v*) − *ε*_*R*_(*v*), is called the CD spectrum. The CD spectrum can be understood in terms of the rotational strength *R*_*if*_ of an electronic transition from an initial state |Ψ_*i*_⟩ to a final state,|Ψ_*f*_⟩, which is defined by the Rosenfeld equation

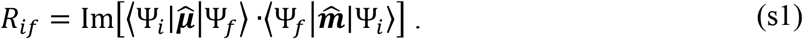

Here 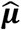 and 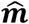 are the electric and magnetic dipole transition moment operators, respectively. The states |Ψ_*i*_⟩ and |Ψ_*f*_⟩ are electronic eigenstates resulting from a chiral arrangement of coupled electric dipole transition moments (or EDTMs), which are each localized to a nucleic acid base residue. Equation (s1) shows that the rotational strength depends on the chirality of the coupled EDTMs, and its sign indicates the handedness (left versus right) of the chiral arrangement.

The Hamiltonian of the coupled system is given by

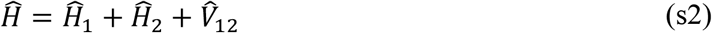

where 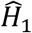 and 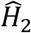 are the Hamiltonian operators of monomers 1 and 2, respectively and 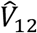 is the coupling between electronic transitions localized to each monomer as defined in the Main Text. The matrix element 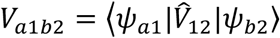 defines the coupling between monomer excited electronic states |*ψ*_*a*1_⟩ (labeled *a* on monomer 1) and |*ψ*_*b*2_⟩ (labeled *b* on monomer 2). The electronic coupling is calculated using the extended-dipole model (EDM),^9^ which has been applied previously to cyanine dyes in self-assembled tubular J-aggregates,^10^ to cyanine dimers in DNA,^11,12^ and to canonical nucleic acid bases in short segments of DNA.^13^ In our current studies, the EDM accounts for the physical length of the adenine base by including for each monomer electronic transition a one-dimensional displacement vector, *l*, that is oriented parallel to the EDTM direction. Each transition dipole moment is represented as two-point charges of equal magnitude and opposite sign (±*q*) separated by distance *l*. The coupling matrix element is given by

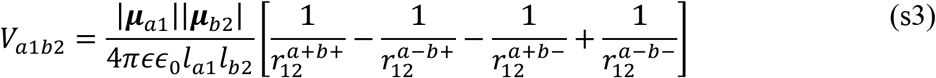

In Eq. (s3), ***μ***_*a*1_ = *q*_*a*1_*l*_*a*1_ and ***μ***_*b*2_ = *q*_*b*2_*l*_*b*2_ are the EDTMs of the transitions a and b on monomers 1 and 2, respectively, and the four distances 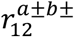 are those between the positive and negative point charges on monomers 1 and 2. The vacuum permittivity of free space is given by *ϵ*_)_, and *ϵ* is the local dielectric constant. For all of our calculations we used the value of the dielectric constant, *ϵ* = 2, in accordance with prior conventions.^14^

In principle, further improvements to the accuracy of our calculations could be achieved by using more detailed, quantum chemical calculations of the electronic transition charge densities. Nevertheless, the favorable comparison between our calculations and experimental data presented below suggests that the EDM provides a reliable estimate of the electronic couplings between adjacent bases for present purposes.

We write the Hamiltonian on a monomer-site basis, such that singly-excited state wave functions are given by tensor products according to

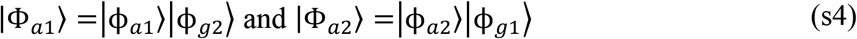

In Eq. (s4), |ϕ_*a*1_⟩ and |ϕ_*a*2_⟩ denote the *a*^th^ electronic excited states of monomers 1 and 2, respectively, and |ϕ_*a*1_⟩ and |ϕ_*a*2_⟩ are the electronic ground states. The number of distinct electronic transitions local to monomer 1 (2) is given by *n*_1(2)_, such that the total number of site-localized transitions is *n*_*tot*_ = *n*_1_ + *n*_2_. The Hamiltonian of Eq. (s2) may thus be written on this site basis as a *n*_*tot*_ × *n*_*tot*_ matrix with diagonal elements representing the single site excitations (with energies *E*_*a*1_ and *E*_*b*2_) and off-diagonal elements representing the couplings *V*_*a*1_*b*_2_ between monomer sites. Note that our formalism neglects the contribution from the isolated adenine monomer, which provides a signal for the fully unstacked conformation. In our calculations, however, this contribution is much smaller than the contribution due to the degenerate coupling of the adenine transitions.

Diagonalization of the Hamiltonian provides the eigen-states |Ψ_2_⟩ and eigen-energies *E*_2_ of the electronically coupled dinucleotide. In the so-called ‘exciton’ basis, the k^th^ singly-excited state |Ψ_2_⟩ may be written

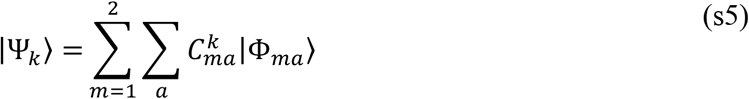

where 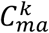 is the expansion coefficient corresponding to transition a local to monomer m. In the exciton basis, the ground state of the dinucleotide is given by |Ψ_*g*_⟩ = |ϕ_*g*1_⟩||ϕ_*g*2_⟩. Using Eq. (s1), we may calculate the rotational strength *R*_*gk*_ (=*R*_*k*_) for the k^th^ electronic transition, where we assign the initial and final states to |Ψ_*g*_ and |Ψ_*k*_⟩, respectively, and the total electric and magnetic dipole transition moment operators are given by vector sums 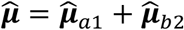 and 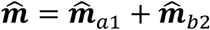.

For a given transition k, the rotational strength depends on the relative orientation of the monomer EDTMs. For the case of coupled degenerate transitions (i.e. *E*_*a*1_ = *E*_*b*2_ and *E*_*k*_ = *E*_*a*1_ + *V*_*a*1*b*2_), the rotational strength is given by

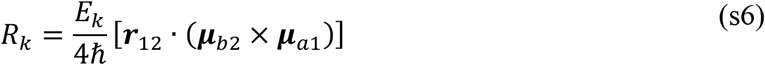

For the case of non-degenerate coupled transitions (i.e. *E*_*a*1_ ≠ *E*_*b*2_ and *E*_2_ ≈ *E*_*k*1_) ≈ *E*_*a*1_, the rotational strength is given by

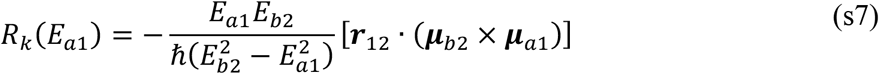

We note that Eq. (s7) is written such that *E*_*a*1_ > *E*_*b*2_.

To calculate the CD spectrum, we consider the relationship between the rotational strength and the integrated area of the CD spectrum within a finite spectral range *v*^5^ →*v*”:

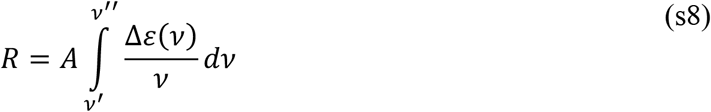

where *A* = 7.659 × 10^−54^ C^2^,^3^s^−1^. For each of the k electronic transitions, we approximate the CD spectral line shape as a Gaussian function 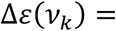 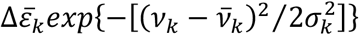, where σ_*k*_ is the Gaussian standard deviation, 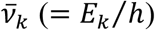 is the mean transition frequency, and 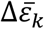 is the magnitude. Upon substitution of the above Gaussian function into Eq. (8), and solving the Gaussian integral, it follows that we may write the magnitude in this approximation as 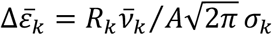. Then, the CD spectral line shape is obtained by summing over all contributions from individual transitions according to

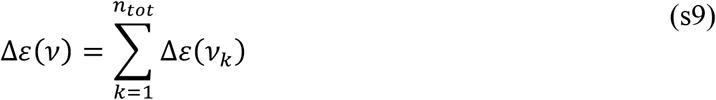

### III. SELECTING THE PARAMETERS FOR THE CALCULATION OF THE CD SPECTRUM

**Figure S2.**
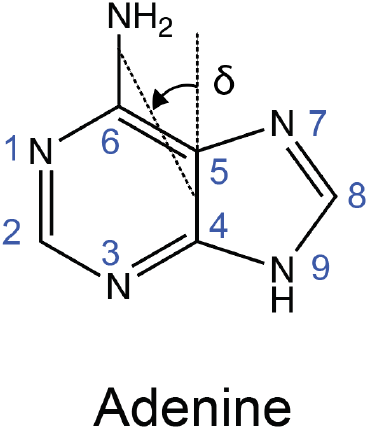
The angle *δ* defines the direction of the electric dipole transition moment (EDTM) used in the CD calculations for the adenine bases of the dApdA dinucleotide monophosphate.

For the majority of our CD calculations, we used as input parameters to Eq. (9) the EDTM data for 9-methyladenine obtained by Holmén et al. (Table S1) ^15^ and the dielectric constant *ϵ* = 2. In Table S1 we list for each transition the values we have used for the EDTM magnitude |*μ*|, orientation *δ*, transition frequency *v*, and extended transition dipole charge *q* and displacement *l* (see Fig. S2). In addition, to model the spectral line width of all monomer electronic transitions we assumed the Gaussian standard deviation σ_*k*_ = 0.2 eV. Our selection of these parameters was based on comparisons between the experimental CD spectrum of dApdA at room temperature in buffer at pH 7.2 containing 0.01 M NaPO_3_ and 0.1 M NaClO_4_, and CD calculations for which we assumed initially that the dApdA dinucleotide adopts only the B-form.

**Table S1.**
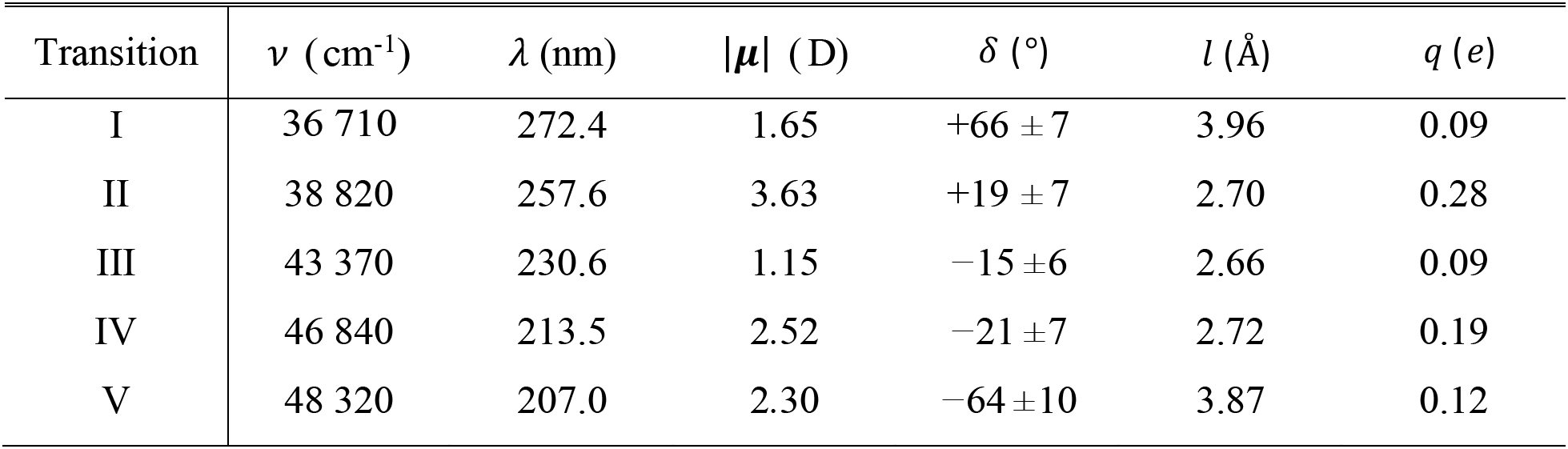
Experimental values for the magnitudes and molecular frame orientations of the electric dipole transition moments (EDTMs) for 9-methyladenine obtained by Holmén et al,^15^ and which we have used to model adenine mononucleotide in this work. All transitions are in-plane *π* → *π*^*^, and are listed in order of increasing transition frequency. The angle *δ* specifies the counter-clockwise rotation of the EDTM vector within the plane of the adenine base relative to the C4-C5 bond axis (see Fig. S2). The partial charges for the extended dipole model were derived using the relation |*μ*| = *q*|*l*|, and by representing the adenine base as an ellipse with major diameter (*a*) 4.6 Å and minor diameter (*b*) 2.6 Å, such that *l* = 2*ab*/[*a*^2^cos^2^(*δ*) + *b*^2^sin^2^(*δ*)]^2^.

For comparison, we present in Table S2 the empirical parameters from Williams et al.^16^

**Table S2.** Empirical spectroscopic parameters from Ref. 16 for the Adenine monomer.

| Transition | $\nu$, cm<sup>-1</sup> | $\mu$, Debye | $\delta$, deg |
| --- | --- | --- | --- |
| I | 37037 | 1.1 | -87 |
| II | 38022 | 4.0 | -3 |
| III | 42553 | 1.0 | -87 |
| IV | 46296 | 3.7 | -87 |
| V | 51282 | 3.7 | -3 |
| VI | 53476 | 4.2 | -87 |

For all of the parameters that we tested (see Tables S1 and S2), we obtained moderately favorable agreement between experiment and theory. We note that the sensitivity of the calculated CD to the choice of input parameters was greatest at the shortest wavelengths (200 – 250 nm) and least at the longer wavelengths (250 – 300 nm).

**Figure S3.**
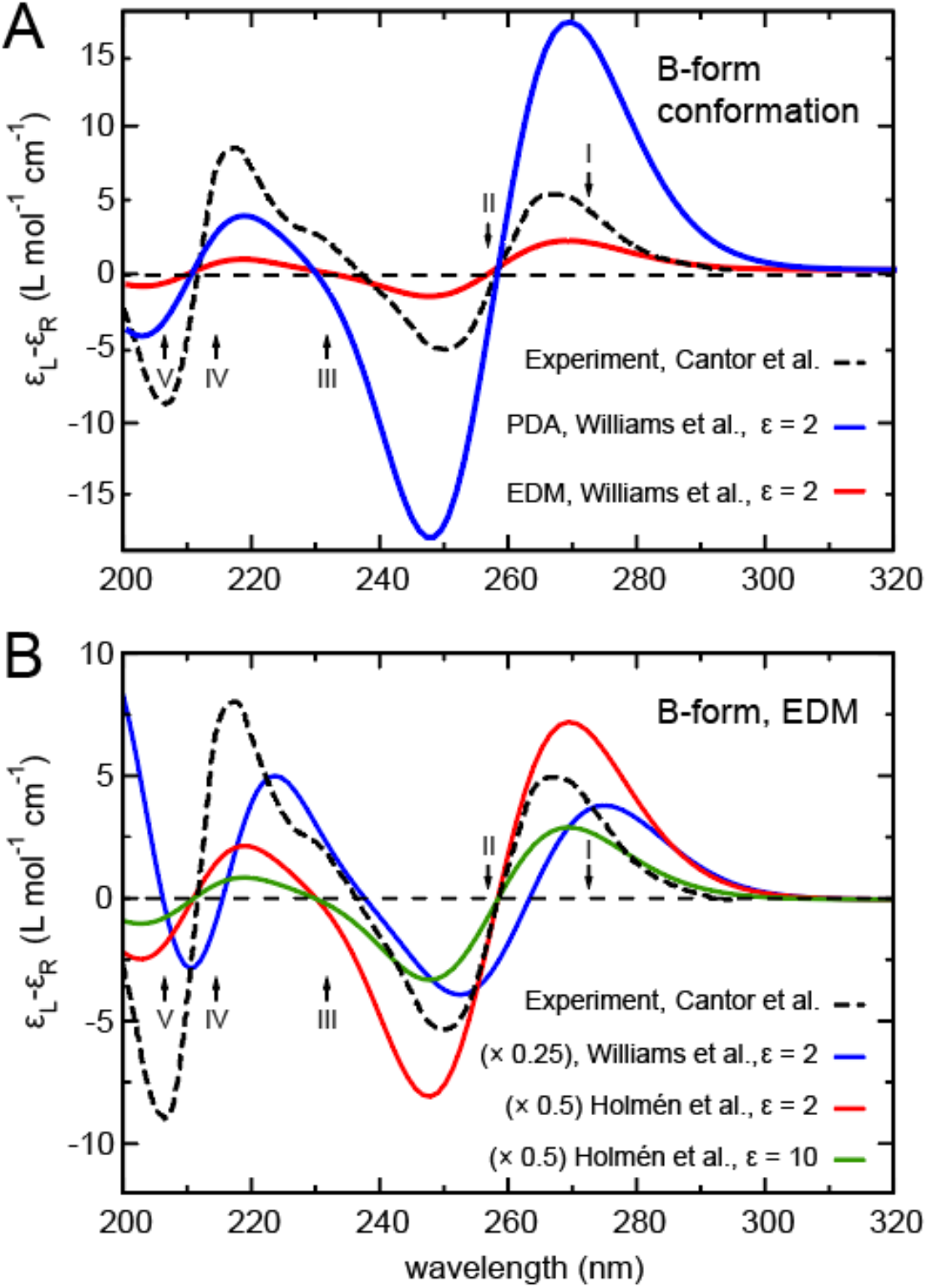
(***A***) comparison of the CD spectrum theoretically predicted for the Watson-Crick B-form of dApdA and the experimental data by Cantor at al.^17^ We show both the spectra calculated using the Point Dipole Approximation (PDA) and the Extended Dipole Model (EDM). (***B***) Comparison of the CD spectrum theoretically predicted for the Watson-Crick B-form of dApdA and the experimental data, using either the empirical parameters from Holmén et al. (Ref. 15) or from Williams et al. (Ref. 16). The effect of varying the dielectric constant (from ϵ = 2 to ϵ = 10) is also shown. In both panels, vertical arrows indicate the positions of the uncoupled transitions of the Adenine monomer listed in Table S1.

To demonstrate the sensitivity of the CD theoretical predictions to the choice of the empirical parameters selected in the CD modeling, we report first, in Fig. S3*A*, a study of the CD spectrum for the Watson-Crick B-form of dApdA calculated using two different models: (i) the simple Point Dipole Approximation (PDA); and (ii) the Extended Dipole Model (EDM). The spectrum of the B-form, predicted by the theory is similar in both approximations, and shows a good agreement with experiments in the low energy part of the spectrum. Figure S3*B* shows, instead, a study of the sensitivity of the calculations to the choice of the parameters. It reports results for the Point Dipole Approximation (PDA) calculation of the CD spectrum for the Watson-Crick B-form, while adopting either the empirical spectroscopic parameters from Holmén et al.^15^ or those from Williams et al.^16^ As can be seen, the positions and heights of the peaks change significantly at low wavelengths (high excitation energies), while they agree reasonably well at high wavelengths (low excitation energies). Thus, the results are qualitatively consistent with a slightly better agreement with the experimental values when using parameters from Table S1. Finally, the figure shows a study of the variation of the empirical dielectric constant, which is found to produce just a change in the intensity of the spectrum, without affecting the positions of the peaks.

### IV. DECOMPOSITION OF THE CD SPECTRUM OF THE dApdA DINUCLEOTIDE MONOPHOSOPHATE

Figure S4 displays the contributions to the CD spectrum of dApdA from each MSM macrostate. We assume only single-excited electronic states and five electronic transitions per adenine monomer, following the conventions used by Holmén et al (ref. 15, Table S1). Proceeding on this basis we obtain ten total degenerate-pair contributions, given by five symmetric and five anti-symmetric transitions. To keep things simple we neglect here the contributions to the spectral decomposition from non-degenerate transitions, because degenerate contributions are dominant in the total CD spectrum (note that both degenerate and non-degenerate contributions are included in the CD calculations reported in the Main Text).

In Fig. S4, different colors represent different transitions, while the dotted and the dashed curves represent the contributions from the symmetric and the anti-symmetric transitions, respectively. Due to the chirality, or lack of chirality, of the average structure in each macrostate, only the states S3 and S4 provide significant contributions to the final CD spectrum. Of particular interest is the spectral decomposition of the two chiral macrostates with the largest stationary probabilities, S3 and S4. The first macrostate includes in its conformational distribution the Watson-Crick B-form, while the second includes the Hoogsteen form.

**Figure S4.**
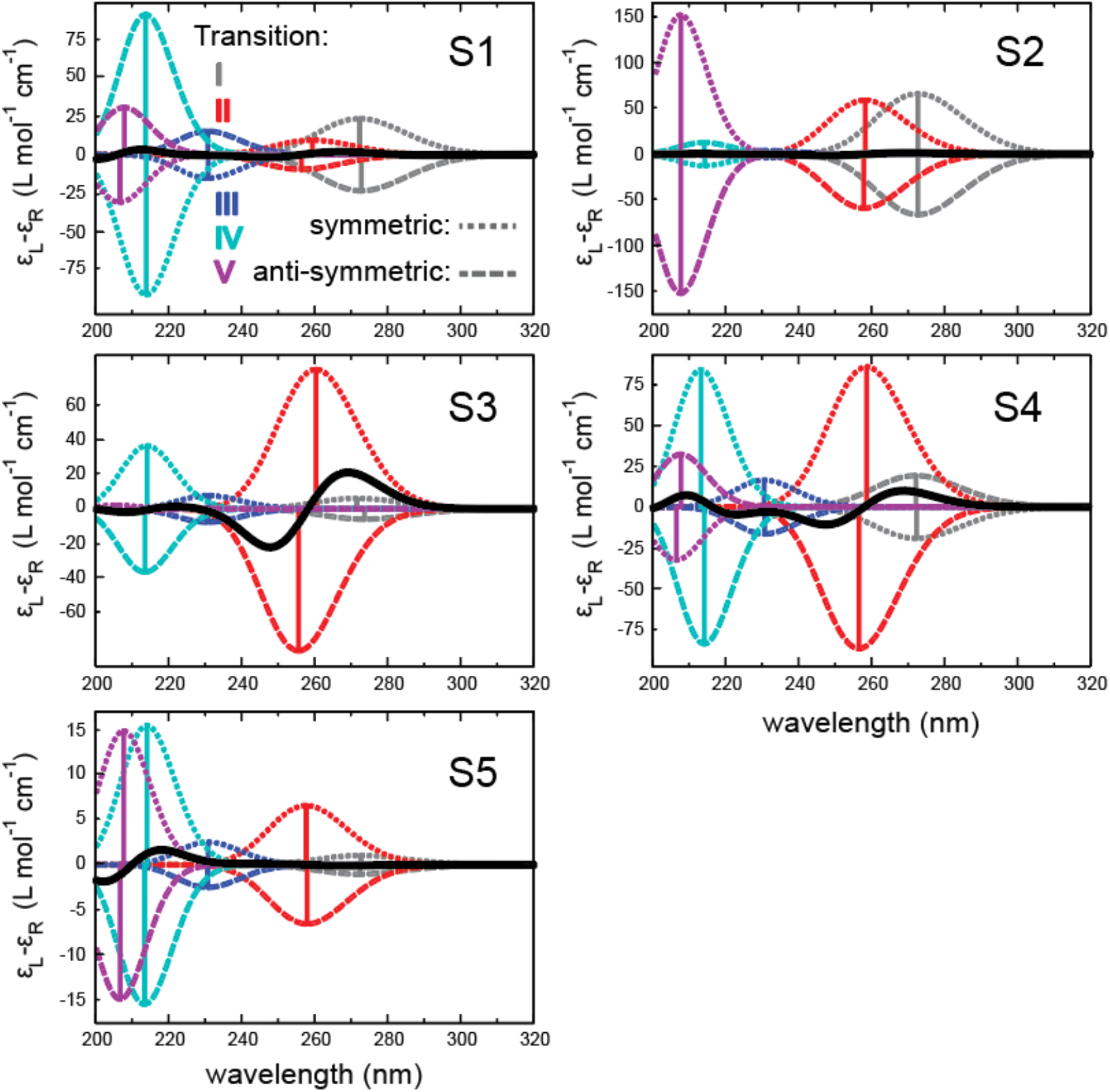
Spectral decomposition of the degenerate CD for the average structures of each of the five macrostates (S1 – S5) for dApdA. The contributions due to transition I (gray), II (red), III (blue), IV (cyan) and V (magenta) are decomposed into signals reflecting the symmetric (dotted) and anti-symmetric (dashed) transitions. The total CD spectrum for each average structure is given by the solid (and thick) black line in each plot.

The average structures of each of these macrostates show that for the dominant low-energy transition (II, red), the decomposition into symmetric and anti-symmetric contributions are of the same sign for both the S3 and the S4 states, while the contributions to the first low-energy transition (I, black) have opposite signs in the two macrostates. The average S3 and S4 macrostate structures, while of opposite handedness, are not mirror images of each other; rather, one of the adenine monomers in the average structure of S4 is ‘flipped’ relative to the same monomer in the average structure of S3, which is compatible with the Hoogsteen structure. It is the flipped base in the S3 macrostate that is responsible for its right-handed CD signal in the long wavelength region.

### V. MODELING THE BASE STACKING OF dApdA USING MARKOV STATE MODELING PROCEDURES

To analyze the base stacking of dApdA, we adopted a two-site description for each base, which defines vectors within the plane of each nucleotide. The first site is positioned between the *C*_4_ and the *C*_5_ carbon atoms in Adenine (see Fig. S5), while the second site was positioned 0.1 nm along a line perpendicular to the vector connecting those atoms.

**Figure S5.**
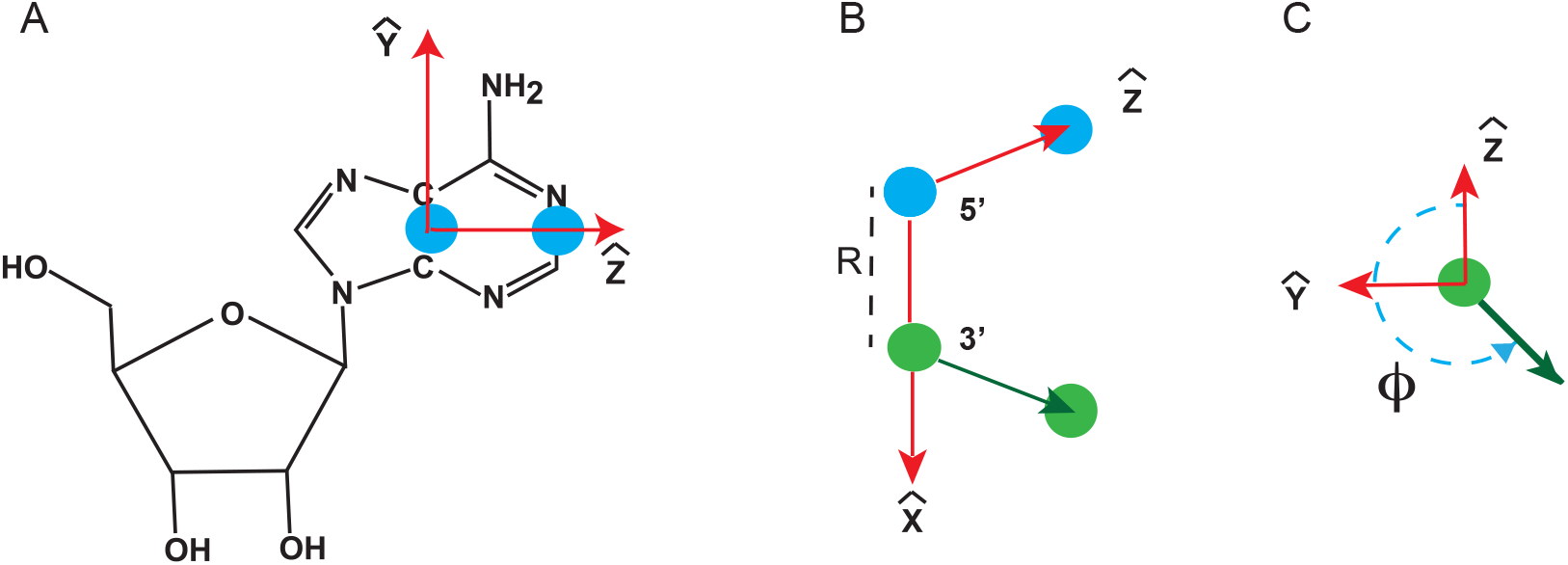
(***A***) The two sites are placed within the plane of the Adenine base. The independent free energy parameters used in the MSM analysis are the radial separation between *C*_4_ and the *C*_5_ midpoints within each adenine monomer, and (***B***) the dihedral angle, *φ*, between the in-plane vectors, here represented in an overhead view (***C***).

The relative orientation of the four-bead model captures the relative distance between the adenine bases, and their relative torsion angle. The parameters reported in the Main Text for the free energy maps are thus defined in Fig. S5: the stacking torsional angle, *φ*, is defined by the dihedral between the in-plane vectors, while the distance, *R*, between nucleotides is given by the distance between the *C*_4_ and the *C*_5_ midpoints of each base (see also Fig. 1 in the Main Text).

### VI. PARTITIONING THE FREE ENERGY SURFACE IN MACROSTATES USING THE MARKOV STATE MODEL PROCEDURE

Briefly, we used the k-means++ algorithm^18,19^ to construct a kinetically-relevant, balanced clustering of the trajectories (using the Euclidean criterion) by partitioning the 10^7^ conformations into 100 initial microstates. A transition rate matrix was constructed for these microstates and then diagonalized into eigenvalues and eigenvectors. From the eigen-spectra of the transition probability matrix, we constructed five macrostates by implementing a minimum error propagation version of the Perron-cluster cluster analysis (PCCA+). We justified our choice for these five macrostates by considering the related conformational landscape and the implied interconversion time scales.

Rapidly interconverting molecular conformations were assigned to the same macrostate, while slowly interconverting conformations, which are separated by large barriers, occur between conformations that lie within different macrostates. By identifying and separating slowly interconverting conformations from rapidly interconverting ones, the MSM ensures that the slow processes obey Markovian statistics. To sample slow transitions, we adopted a lag-time of 500 ps, and confirmed that under these conditions Markovian behavior was satisfied by checking that the Chapman-Kolmogorov condition applies^20,21,22^(see Fig. S6).

We started by assigning the simulation trajectory to W=100 discrete microstates using the k-means++ clustering algorithm as implemented in PyEMMA^23^ software program (the comparison with the case of assuming W=1000 is reported at the end of this section and shows no substantial difference in their predictions). Once the clustering of the simulation trajectory into microstates was completed, we defined a lag time τ and calculated the transition matrix, *T*(τ), by counting the number of transitions occurring between two given microstates during the defined lag time. The transition matrix, therefore, models the evolution of the probability vector, *P*^*T*^(*t* + τ) = *P*^*T*^(*t*)*T*(τ), which gives the probability of finding the system in a final state at time t+τ, given that the system was in an initial state at time t, and had a probability of transition between states, during lag time τ, that is given by the transition matrix, *T*(τ).

The diagonalization of the transition matrix gives eigenvalues and eigenvectors, which contain important information on the stable states in the free energy landscape and the kinetics of transitions between states. The eigenvalues (λ) define the timescale of a transition *i* during the lag time τ as *t*_*i*_ = −τ/[ln *λ*_*i*_(τ)], while the eigenvectors define the partitioning of the free energy surface, and its 100 microstates, into a smaller number of macrostates. Because the transition matrix is a regular stochastic matrix, the Perron-Frobenius theorem guarantees that the first eigenvalue is equal to one, which corresponds to an infinite transition time *t*_1_. Thus, the corresponding first eigenvector has only positive entries, and defines the equilibrium (*t*_1_ = ∞) population of the macrostates, which in our calculation of the CD spectrum gives the percentage contribution of each macrostate to the final CD function (see Fig. 6 in the Main Text). By inspecting the eigenvalues, it is possible to identify a gap, corresponding to a gap in the transition times, which separates slow from fast transitions. The eigenvector that corresponds to that eigenvalue defines, with the number of its “nodes”, the number of macrostates into which the free energy landscape needs to be partitioned to separate fast transitions (inside one macrostate) from slow, and biologically relevant, transitions (between macrostates). In our case this condition is fulfilled when the free energy map is partitioned into five macrostates (see Fig. 5*A* in the Main Text). The lag time, τ, at which the kinetics of the stacking-unstacking fluctuations become uncorrelated, is identified by testing at which τ the transitions between states become Markovian. To test the Markovian nature of those transitions, we adopted the standard procedure based on fulfilling the Chapman-Kolmogorov condition. When dynamical processes are Markovian (i.e. uncorrelated), the transition matrix sampled at a multiple, *n*, of the lag time, τ, is equal to the transition matrix at lag time τ to the *n* power: *T*(*n*τ) = *T*(τ)^7^, which implies that the eigenvalues *λ*(*n*τ) = *λ*(τ)^7^. As a consequence, the timescale of a transition becomes independent of the time used to sample the simulation trajectory. In fact 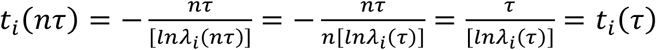, which is the Chapman-Kolmogorov condition.

To test this condition, Fig. S6 reports the transition time as a function of the lag time and identifies the time at which the process becomes Markovian as the time where *t*_#_(*n*τ) becomes constant. Figure S6 shows that in our system there are four slow processes that are fully Markovian when transitions are sampled at a lag time longer than 500 ps. Thus, a lag time of 500 ps has been selected for the calculations reported in the Main Text. The number of slow Markovian processes detected in this plot, which is defined as the number of lines that becomes constant while fulfilling the necessary condition that *t*_*i*_(τ) ≥ τ, is consistent with a number of macrostates equal or higher than five, thus supporting the number of macrostates selected in our analysis.

As can be seen in Fig. S6, beyond 0.5 ns, the transition time (implied timescale) is almost level. This means that starting at 0.5 ns, and at longer time lag, the coarse graining of the free energy map into five macrostates gives kinetics transitions between macrostates that are Markovian, indicating that a reasonable separation of timescales exists in the spectral decomposition of the transition matrix. The fulfillment of the Chapman-Kolmogorov condition ensures that the transition time is independent of the number of uncorrelated steps that are used to model the process.

**Figure S6.**
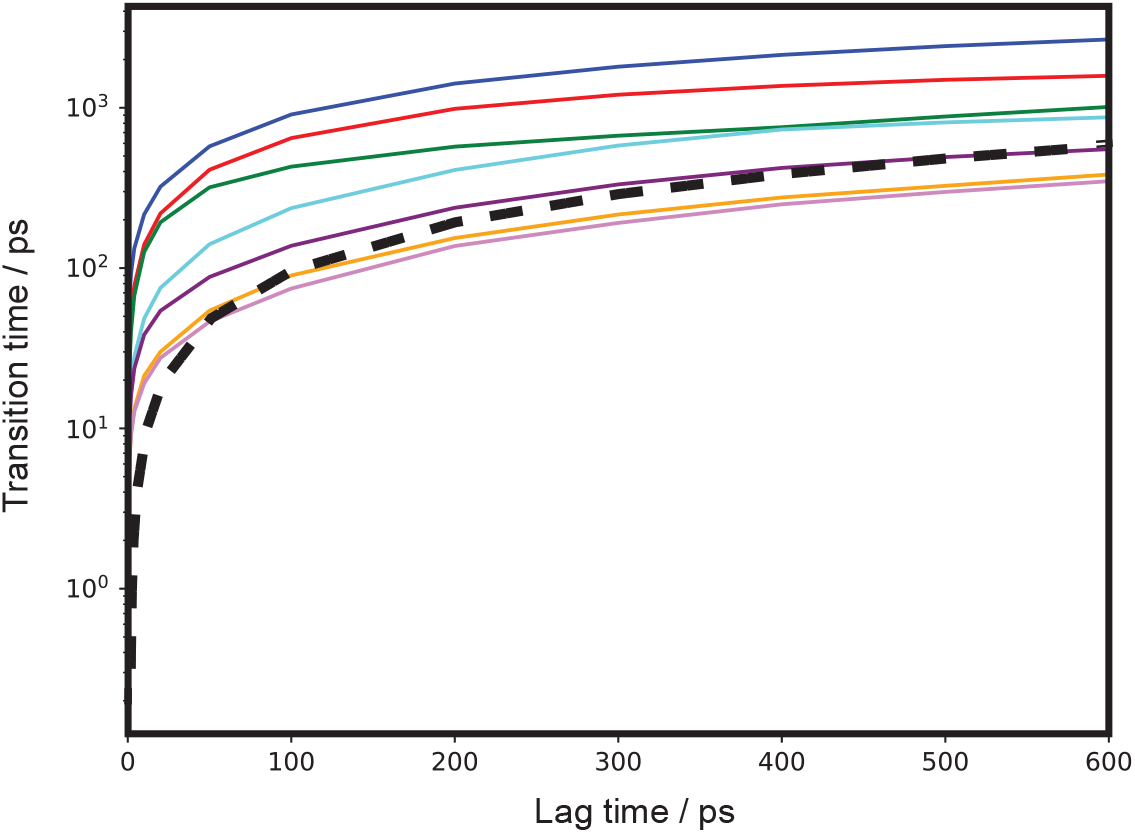
Transition time measured as a function of increasing lag time, for the simulation of dApdA in solution at 0.1 M salt concentration, where the free energy landscape is partitioned into five Markov states. Different colors display different transition processes. For the four slowest processes the transitions become Markovian around 500 ps. The black dashed line defines the condition for which the transition time is equal to the sampling lag time: processes that occur in a time faster than the sampling time cannot be sampled and are discarded.

### VII IDENTIFYING THE OPTIMAL NUMBER OF MICROSTATES

The number of microstates to be used in the MSM analysis depends on the precision we want to achieve in the MSM calculations, while avoiding underfitting and overfitting. The optimal number is roughly related to the number of structural parameters and the number of groups/residues in the molecule. Here, we studied a relatively small system (dApdA) using simple structural parameters, which suggests that a small number of microstates could be sufficient. Note that when the number of microstates is not optimum, the kinetic information and/or the positions of the barriers and the border between metastable states can be inaccurately predicted because of underfitting or overfitting. In this Section, we investigate the sensitivity of the CD spectra predictions and the related kinetic information to the number of microstates and compare the results for 100 and 1000 microstates, while in the Main Text we reported the results for 100 microstates.

**Table S3.** Stationary normalized probability distributions of PCCA+ macrostates when the MSM contains either 100 or 1000 microstates.

|  | S1 | S2 | S3 | S4 | S5 |
| --- | --- | --- | --- | --- | --- |
| 100 microstates | 0.026 | 0.028 | 0.090 | 0.373 | 0.483 |
| 1000 microstates | 0.024 | 0.027 | 0.095 | 0.372 | 0.482 |

Table S3 reports the stationary distribution (i.e. the first eigenvector of the transition matrix) for each of the macrostates in the 100- and 1000-microstate MSM models (see also Fig. 6 in the Main Text). As it can be seen, there is no significant difference between them, indicating that selecting either number of microstates does not entail significant changes in the borders between macrostates. Thus, the slight change in border resolution does not significantly affect the long-time properties of the Markov chain, whether 100 or 1000 microstates were used.

**Table S4.** Mean first passage times (MFPTs) in nanoseconds between the five macrostates of the dApdA system at [NaCl] = 0.1 M. The MFPTs were calculated using the Markov state model analysis of the free energy landscape (Fig. 5*A* in the Main text) and 100 microstates. Note that the unit of time is ns and rows and columns indicate initial and final macrostates, respectively.

| $i \setminus f$ | S1 | S2 | S3 | S4 | S5 |
| --- | --- | --- | --- | --- | --- |
| S1 | 0.0 | 35.0 | 13.1 | 5.9 | 3.2 |
| S2 | 56.4 | 0.0 | 12.6 | 4.5 | 3.2 |
| S3 | 58.7 | 37.0 | 0.0 | 6.1 | 2.4 |
| S4 | 59.1 | 36.4 | 13.4 | 0.0 | 4.6 |
| S5 | 58.1 | 36.9 | 11.5 | 6.5 | 0.0 |

In Table S4 we show the mean first passage times (MFPTs) between the macrostates when 100 microstates are used to build the transition matrix for the MSM. When compared to the analogous table for 1000 (see Table S5), it shows that there is no significant difference between the MFPTs calculated using 100 or 1000 microstates. Finally, we compare the behavior of the implied timescales as a function of the lag time for the 100- and the 1000-microstate MSMs, and we find that the transition time fulfills the Chapman-Kolmogorov condition at a lag time consistent in the two cases (data not shown).

**Table S5.** Mean first passage times (MFPTs) in nanoseconds between the five macrostates of the dApdA system at [NaCl] = 0.1 M. The MFPTs were calculated using the Markov state model analysis of the free energy landscape (Fig. 2*A* in the Main text) and 1000 microstates. Note that the unit of time is ns and the initial and target macrostates are in rows and columns, respectively.

| $i \setminus f$ | S1 | S2 | S3 | S4 | S5 |
| --- | --- | --- | --- | --- | --- |
| S1 | 0.0 | 36.2 | 13.0 | 6.3 | 2.9 |
| S2 | 68.4 | 0.0 | 12.3 | 4.6 | 3.3 |
| S3 | 70.9 | 38.3 | 0.0 | 6.2 | 2.5 |
| S4 | 71.5 | 37.6 | 13.1 | 0.0 | 4.7 |
| S5 | 69.9 | 38.2 | 11.2 | 6.6 | 0.0 |

### VIII SIMPLIFIED CD SPECTRAL CALCULATION USING MSM AVERAGED MACROSTATE CONFORMATIONS

In the Main Text, we showed that the CD spectrum of the dApdA system can be decomposed into contributions from five different MSM macrostates, which are defined based on their bounded positions within the free energy landscape *G*(*R*, *φ*) of 10 million MD-sampled microstate configurations (see Fig. 2*D*). Of the five macrostates, only S3 and S4 contribute significantly to the CD spectrum (see Fig. 6*F*). Our results suggest that we may apply a relatively simple model to achieve the structural interpretation of the dApdA CD spectrum. Having identified the key macrostates relevant to the CD observable (see above), we next determine the smallest number of structural parameters necessary to characterize these macrostates. Given a reduced set of conformations for the various macrostates, in addition to specification of their relative weights, it is possible to greatly speed up the computation time needed to simulate the CD spectrum. In principle, such structural models can be used for the general interpretation of any spectroscopic measurement performed on the dApdA system.

For each of the five macrostates we determined an average conformation with mean and standard deviation of the inter-base separation defined according to

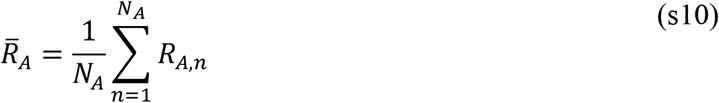

and

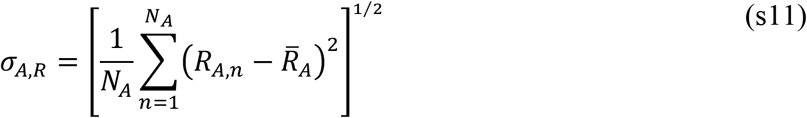

We similarly defined the means and standard deviations of the inter-base twist, tilt and roll angles according to

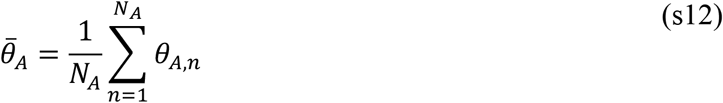

and

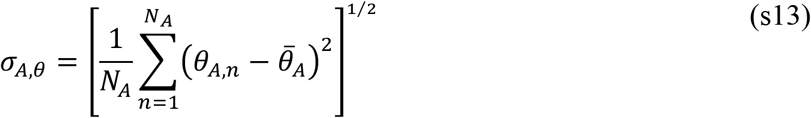

where 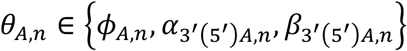 are the *n*th twist, tilt and roll angles, respectively, of the *N*_*A*_ configurations contained within the *A*th macrostate (see Fig. 1 in the Main Text and Fig. S5 for coordinate definitions). In Table S6, we list the mean and standard deviation parameters for each macrostate, in addition to the associated number fractions *p*_*A*_ (= *N*_*A*_/*N*_*tot*_) calculated from the MSM analysis.

In Figs. S7*A* – S7*E* we show our CD calculations for macrostates S1 – S5, respectively, which were determined from the mean macrostate conformations (shown as red curves). We compare these to the calculated CD spectra by summing over all of the configurations contained within each macrostate (blue curves).

**Table S6.** Means and standard deviations of the structural parameters corresponding to the five macrostates *A*, for dApdA, which are labeled S1 – S5. The mean inter-base separation 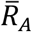 mean twist angle 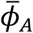, mean tilt angles 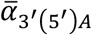, and roll angles 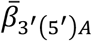 are defined by Eqs. (s10) – (s13), which are based on the structural coordinates defined in Fig. 1 of the Main Text and Fig. S5. The values for the equilibrium population, *p*_A_, are used as weights for computing the CD spectrum from the mean macrostate conformations, as shown in Fig. S7.

| $A$ | $p_A$ | $\bar{R}_A$<br>(Å) | $\sigma_{A,R}$<br>(Å) | $\bar{\phi}_A$<br>(°) | $\sigma_{A,\phi}$<br>(°) | $\bar{\alpha}_{3'(5')A}$<br>(°) | $\sigma_{A,\alpha_{3'(5')}}$<br>(°) | $\bar{\beta}_{3'(5')A}$<br>(°) | $\sigma_{A,\beta_{3'(5')}}$<br>(°) |
| --- | --- | --- | --- | --- | --- | --- | --- | --- | --- |
| S1 | 0.026 | 4.5 | 1.1 | 145.0 | 6.0 | 155.3<br>(28.6) | 9.8<br>(10.6) | 33.2<br>(-28.9) | 39.7<br>(51.4) |
| S2 | 0.028 | 12.2 | 1.4 | 12.0 | 183.6 | 160.1<br>(23.8) | 5.3<br>(6.3) | -58.6<br>(-4.1) | 61.9<br>(59.8) |
| S3 | 0.090 | 4.0 | 0.3 | 44.0 | 1.8 | 139.0<br>(43.5) | 8.3<br>(8.6) | -28.6<br>(-14.3) | 82.2<br>(72.2) |
| S4 | 0.373 | 4.7 | 0.9 | -91.7 | 15.2 | 148.5<br>(33.2) | 9.5<br>(9.8) | -76.2<br>(21.3) | 28.9<br>(56.1) |
| S5 | 0.483 | 5.0 | 0.9 | 5.9 | 10.0 | 159.0<br>(22.7) | 7.4<br>(7.6) | -69.3<br>(32.5) | 29.4<br>(38.9) |

The insets in Figs. S7*A* – S7*E* show the molecular models of the dApdA dinucleotide that represent the corresponding mean macrostate conformations. For macrostates S1, and S3 – S5, the calculated CD spectrum of the mean conformation are similar in shape to those of the full macrostates, although the magnitude of the CD in each case is somewhat overestimated by the mean conformation. This agreement is less favorable for macrostates S2, which is likely due to the broad dispersion of conformations contained within this macrostate.

**Figure S7.**
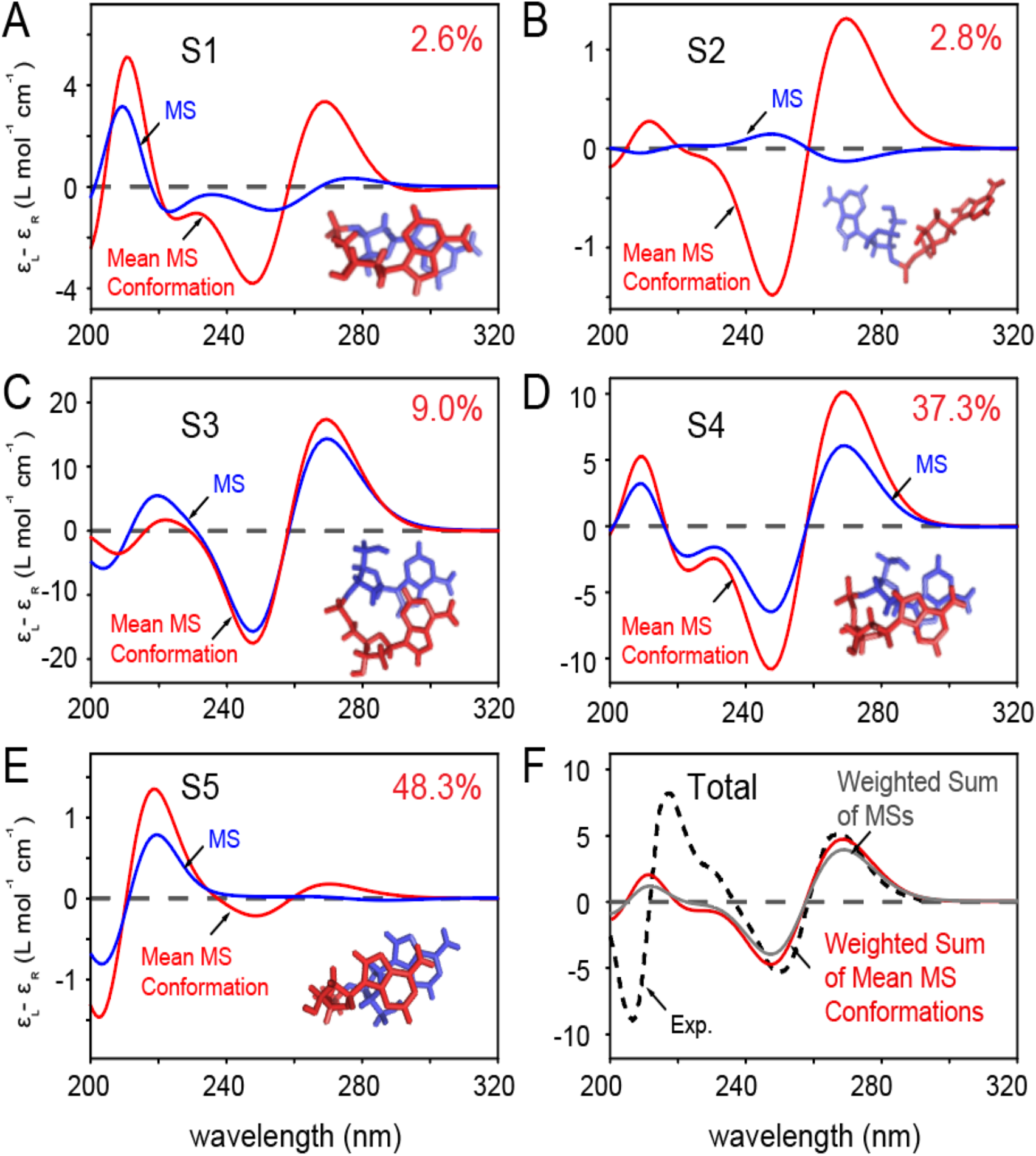
(***A***) – (***E***) Each panel shows, for each macrostate, the comparison between the contribution to the CD spectrum from all the conformational states in the macrostate (blue curve) and the contribution from the averaged macrostate structure (red curve), with structural parameters listed in Table S6. The molecular models representative of the averaged dApdA structures are shown as insets within each panel. The 5’ nucleotide is shown in blue, and the 3’ nucleotide and phosphate are shown in red. (***F***) The weighted sum of the macrostate contributions to the total CD are shown in gray, and the weighted sum from the averaged structures in red. The experimental CD spectrum^7,12^ is shown as a dashed black curve.

As discussed in the Main Text, the two macrostates that contribute most significantly to the CD spectrum are S3 and S4. Our determination of the mean conformations allows us to apply a structural interpretation to these contributions. Macrostate S3 exhibits, on average, a stacked and right-handed conformation, while macrostate S4 exhibits a stacked and left-handed conformation. The S3 mean conformation resembles that of flanking bases in B-form duplex DNA, which gives rise to the ‘right-handed’ Cotton effect observed in the long wavelength region of the CD spectrum. The relative roll angle of the S4 conformation is large 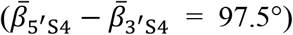, such that the 3’ base is ‘flipped’ relative to the 5’ base. This conformation also gives rise to the right-handed long wavelength CD spectrum. A spectral decomposition analysis of macrostate S4 shows that the right-handed long wavelength CD is consistent with this flipped base conformation (see Fig. S3 above). Although the S5 macrostate represents 48.3% of the total population, it contributes very little to the CD spectrum due to its predominantly achiral symmetry and correspondingly low rotational strength. Finally, the S1 and S2 macrostates do not contribute significantly to the CD spectrum due to their small populations.

### IX DISTRIBUTIONS OF THE THETA ANGLE AT INCREASING WATER SEPARATION FROM THE PHOSPATE OXYGEN FOR dApdA

In Fig. S8, we report the distributions of the angle *θ* that defines the orientation of the water dipole moment 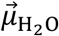 relative to the vector 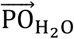, which connects the water oxygen to the central phosphorous atom P of the dApdA (see Fig. 4*A* of the Main Text). The distributions are reported for the dApdA dinucleotide at monovalent salt concentration [NaCl] = 0.1 M. Each distribution is defined over a narrow range of distances corresponding to a given hydration shell relative to the P atom. With the exception of the first distribution, which displays an irregular structure due to the exclusion of water molecules from the nearest distances around the P atom, all of the remaining distributions appear as continuous and smoothly varying functions of the angle *θ*. The distributions exhibit a symmetric shape only for the hydration layers that are separated from the P atom by more than 7 Å. For shorter distances, we observe well-defined non-uniform distributions of the water dipole orientations, which indicate the presence of hydrogen bonding between successive hydration shells. From our calculations of the average angle, we see that at short distances < 〈cos *θ*〉 ≈ cos〈*θ*〉 as shown in Fig. 4 of the Main Text.

**Figure S8.**
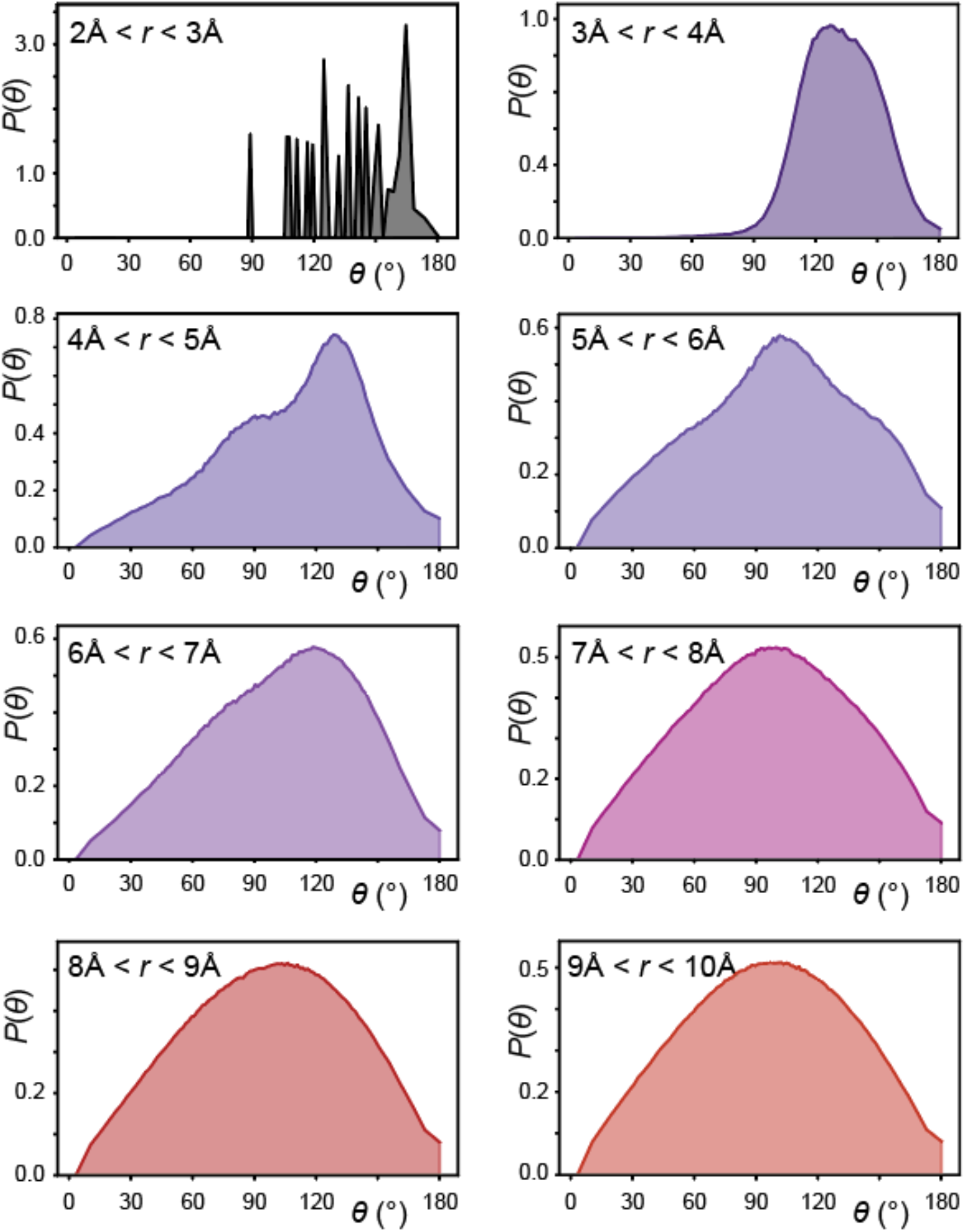
Distributions of the angle *θ* between the water dipole moment 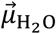 and the vector 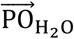, as a function of distance between the water oxygen and the central phosphorous atom P. The distributions are shown as a function of increasing distance from the P atom, and for the salt concentration [NaCl] = 0.1 M. The non-uniform broadening of the distributions indicate the presence of hydrogen bonding between successive hydration shells and loss of orientational correlation with the P atom with increasing distance.

### X DECAY OF THE TIME AUTOCORRELATION FUNCTION OF THE INTER-BASE SEPARATION, R(t)

To ensure that the system is converged and that the FES displayed in Fig. 2 in the main text represents the system at equilibrium, we report in Fig. S9 the decay of the autocorrelation function of the fluctuations away from the equilibrium value of the inter-base distance, Δ*R*(*t*) = *R*(*t*) − 〈*R*〉.

The autocorrelation function is defined 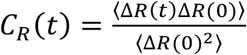. The autocorrelation function decays to zero, for all the salt concentrations, on a timescale of a few nanoseconds. Note that the decay is similar at all salt concentrations up to [NaCl]=1. M, but becomes faster in the high salt concentration regime ([NaCl]=1.5 M), where bases are largely unstacked.

**Figure S9.**
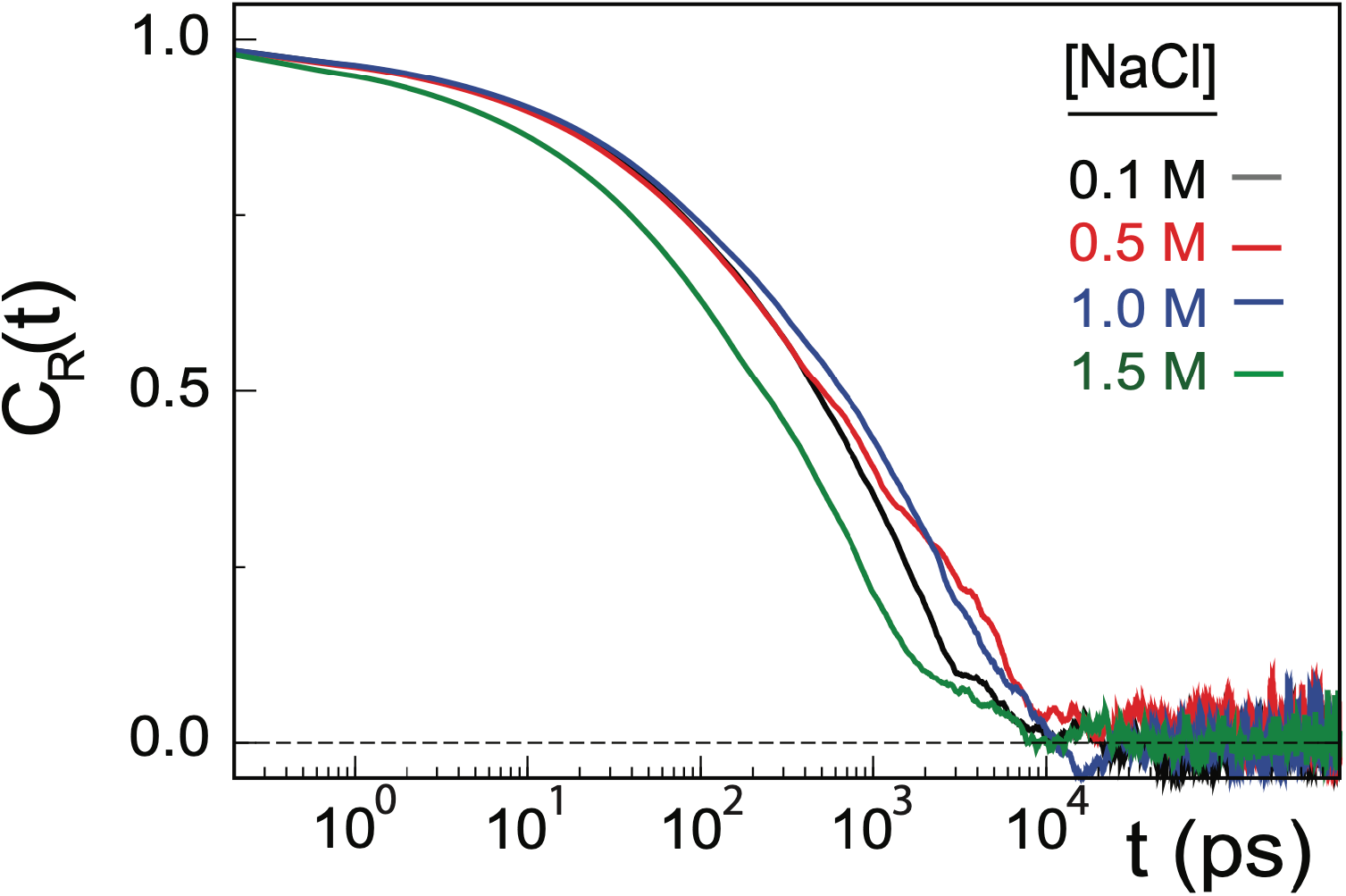
Decay of the time autocorrelation function of the fluctuations of inter-base separation for samples at increasing salt concentration. The system reaches equilibrium in approximately 10 ns, with the sample at salt concentration higher than 1.0 M relaxing faster than any of the other salt concentrations. In contrast, the samples at salt concentrations below 1 M decay at very similar rates.

